# Mitogenomes illuminate the origin and migration patterns of the indigenous people of the Canary Islands

**DOI:** 10.1101/486142

**Authors:** Rosa Fregel, Alejandra C. Ordóñez, Jonathan Santana-Cabrera, Vicente M. Cabrera, Javier Velasco-Vazquez, Verónica Alberto, Marco A. Moreno-Benítez, Teresa Delgado-Darias, Amelia del Carmen Rodríguez-Rodríguez, Juan Carlos Hernández, Jorge Pais, Rafaela González-Montelongo, José M. Lorenzo-Salazar, Carlos Flores, M. Carmen Cruz de Mercadal, Nuria Álvarez-Rodríguez, Beth Shapiro, Matilde Arnay, Carlos D. Bustamante

## Abstract

The Canary Islands’ indigenous people have been the subject of substantial archaeological, anthropological, linguistic and genetic research pointing to a most probable North African Berber source. However, neither agreement about the exact point of origin nor a model for the indigenous colonization of the islands has been established. To shed light on these questions, we analyzed 48 ancient mitogenomes from 25 archaeological sites from the seven main islands. Most lineages observed in the ancient samples have a Mediterranean distribution, and belong to lineages associated with the Neolithic expansion in the Near East and Europe (T2c, J2a, X3a…). This phylogeographic analysis of Canarian indigenous mitogenomes, the first of its kind, shows that some lineages are restricted to Central North Africa (H1cf, J2a2d and T2c1d3), while others have a wider distribution, including both West and Central North Africa, and, in some cases, Europe and the Near East (U6a1a1, U6a7a1, U6b, X3a, U6c1). In addition, we identify four new Canarian-specific lineages (H1e1a9, H4a1e, J2a2d1a and L3b1a12) whose coalescence dates correlate with the estimated time for the colonization of the islands (1^st^ millennia CE). Additionally, we observe an asymmetrical distribution of mtDNA haplogroups in the ancient population, with certain haplogroups appearing more frequently in the islands closer to the continent. This reinforces results based on modern mtDNA and Y-chromosome data, and archaeological evidence suggesting the existence of two distinct migrations. Comparisons between insular populations show that some populations had high genetic diversity, while others were probably affected by genetic drift and/or bottlenecks. In spite of observing interinsular differences in the survival of indigenous lineages, modern populations, with the sole exception of La Gomera, are homogenous across the islands, supporting the theory of extensive human mobility after the European conquest.

## Introduction

The Canaries archipelago is located off the southern coast of Morocco (Figure 1). Due to their oceanic volcanic origin, they have probably never been connected to the continent. Mediterranean sailors discovered several groups of islands in the Atlantic Ocean in the 13^th^ century, but only the Canary Islands were found to be inhabited by an indigenous population [1]. European chroniclers recorded that different islands were inhabited by populations exhibiting different ways of life and speaking distinct dialects of what they believed to be a Berber language. Ethno-historical sources provided ethnonyms for the native population of each island (e.g. Guanches for Tenerife, Benehaoritas for La Palma, and Bimbapes for El Hierro). However, for clarity, we will refer to them in general terms, as the Canarian indigenous or native population.

**Figure 1.**
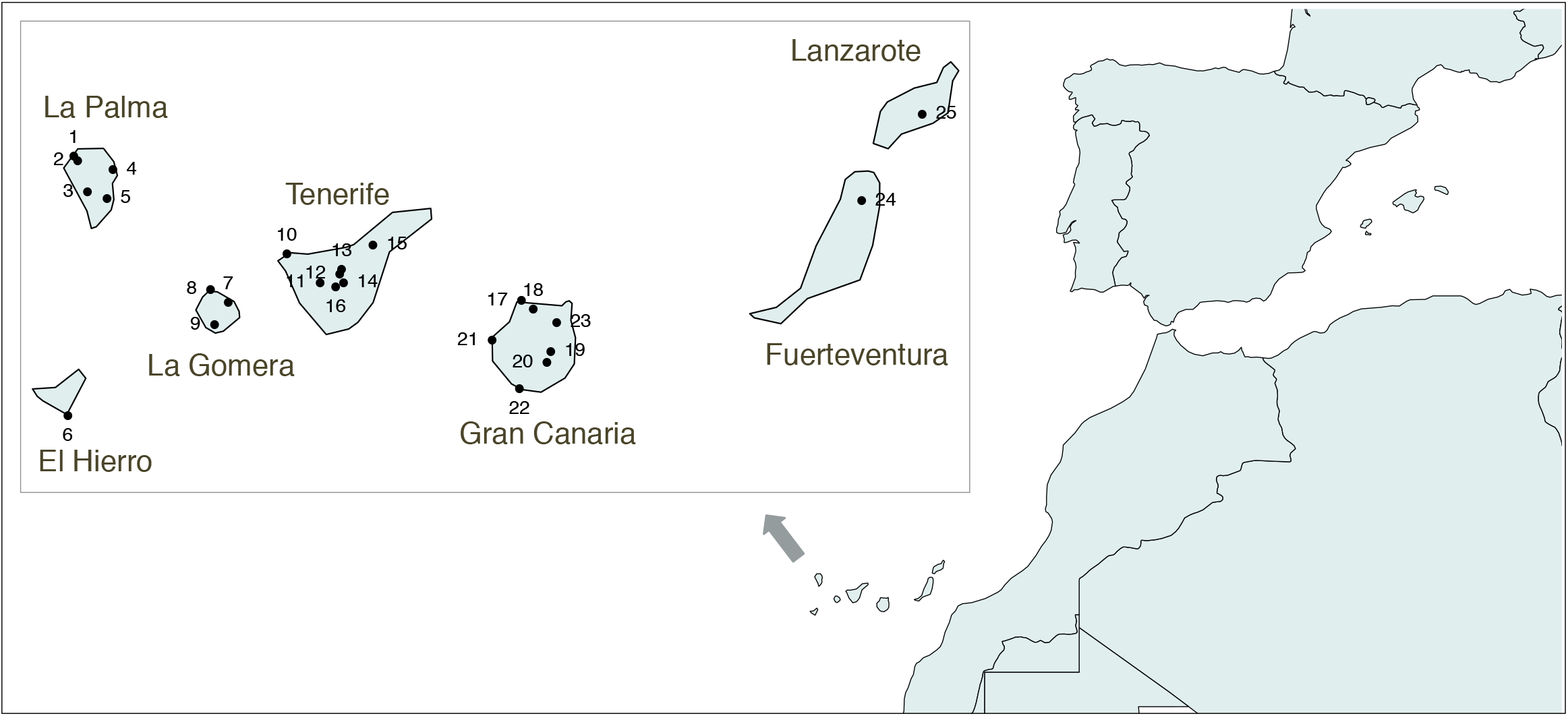
Map of the Canary Islands showing the geographical location of the archaeological sites included in this study. Codes are as follows: 1 – Cueva del Agua; 2 – Huerto de los Morales; 3 – Salto del Casimiro; 4 – El Espigón; 5 – Los Pasitos; 6 – Punta Azul; 7 – Barranco de Majona; 8 – El Pescante; 9 – Antoncojo; 10 – Las Arenas; 11 – El Cedro; 12 – El Salitre; 13 – El Portillo; 14 – La Angostura; 15 – El Cascajo; 16 – El Capricho; 17 – El Agujero; 18 – El Hormiguero; 19 – Guayadeque; 20 – La Fortaleza; 21 – Cuermeja; 22 – Lomo Galeón; 23 – Puente de la Calzada; 24 – El Huriamen; 25 – Montaña Mina.

Chroniclers were amazed to discover that the Canarian natives were unaware of navigational methods and had remained isolated from the African continent [2,3]. During the 15^th^ century, the Spanish kingdom of Castile gradually conquered all of the Canary Islands, after previous European attempts. In most of the islands, the indigenous people resisted the European conquest [4]. The crushing of the resistance, and subsequent European colonization, had a great impact on the indigenous people [5]. In spite of the abolishment of slavery on the Islands in 1498, a large number of natives were deported during and after the conquest [6]. Those that survived and stayed progressively mixed with the European colonizers, leading to the loss of indigenous culture and language.

The geographic origin of the Canarian indigenous people was initially inferred from both the interpretation of historical written sources and the analysis of archaeological evidence. Most archaeological and anthropological data support a North African origin for the Canarian indigenous people, relating to the Berber populations [7,8]. Key evidence supporting a Berber origin includes inscriptions belonging to the Libyco-Berber and Lybico-Canarian alphabets [9,10], pottery [11], communal granaries [12], and domestic species [13–15]. Non-metric dental traits [16–18] and morphological analyses of cranial and long bones [19,20] also show similarities between current inhabitants of Northwest Africa and the Canarian indigenous people.

In regards to the time of the arrival of the first population groups, some authors have proposed the first millennium BCE as the upper bound for human presence in the archipelago [21], based on radiocarbon dating of charcoal and sediment samples. In addition, there is evidence of a Roman short-stay settlement in Lobos islet dated during or before the first centuries of the present era [22], which did not, according to the archaeological data, involve attempting to colonize the Canaries. Recently, there has been an effort to review and contextualize radiocarbon dates in the Canary Islands to better assess the time of the archipelago’s indigenous colonization. Accelerator mass spectrometry (AMS) analyses support a later colonisation of the Canary Islands dating to the outset of the first millennium AD. If only AMS analyses performed on short-lived samples are considered [23], the earliest dates from the eastern islands of Lanzarote and Fuerteventura range between 100–300 cal AD [21,22], whereas those from the central island of Gran Canaria range between 400–500 AD [3]. The oldest AMS dates from Tenerife are around 660–880 cal AD [24], while the western islands of La Palma, El Hierro and La Gomera yield AMS dates ranging respectively between 260–450 cal AD [24], 420–610 cal AD [24], and 120–330 cal AD [25]. On the other hand, older radiocarbon dates that place the arrival of human populations before the 1^st^ century BCE were obtained from sediment, wood and charcoal samples that could be older than the archaeological site where they were excavated.

Mitochondrial DNA (mtDNA) is a powerful tool for inferring the geographic origin of populations [26]. MtDNA is maternally inherited, does not undergo recombination and its different lineages are geographically structured in human populations. For those reasons, mtDNA has been widely applied in phylogeographic studies. The analysis of current Canary Islands samples using mtDNA has provided support for a North African origin for the indigenous people, based on the presence of the mtDNA U6 haplogroup [27], which has a clear Berber ascription [28,29]. Within the U6 lineages observed in the current Canary Islanders, it is worth mentioning U6b1a, a haplogroup that is not present today in North Africa and which is considered a Canarian autochthonous lineage [30]. Interestingly, U6b1a’s coalescence age (3,600 years ago) predates the proposed time of arrival of the first inhabitants of the islands, suggesting an origin in North Africa [30]. Other haplogroups observed in the current Canarian people have Eurasian (H, T, J…), sub-Saharan African (L1, L2 and L3) and Amerindian (A2 and C1) affiliations [31]. These results highlighted the multiethnic nature of the modern population of the Canary Islands, correlating with historical events, such as the implementation of a slave workforce for the sugar cane plantations, or the commercial connection with the Americas in the colonial period [32]. The detailed analysis of current mtDNA of the modern Canary islanders has also suggested possible origins for the indigenous population, including Morocco, Tunisia, Algeria or Sahara, but an overall agreement has not yet been reached [31,33].

Regarding the colonization model, linguistic research has pointed to at least two migration waves from North Africa [10,34]. Also, the observation of different cultural backgrounds affecting the island of La Palma has been interpreted as evidence of consecutive migrations. The specific timing for those migrations is still unclear, except for La Palma, where the second wave of migration has been proposed to have taken place around the 10^th^ century [7]. This idea has also been supported by asymmetrical distribution of both mtDNA [31] and Y-chromosome lineages [35] in the modern Canarian population. The first colonization wave may have affected the entire archipelago, creating the substrate population and bringing mtDNA and Y-chromosome haplogroups observed today in most of the islands, including the mtDNA lineages U6b1a or H1cf. The second colonization would have brought new migrants to certain islands and created an asymmetrical distribution of haplogroups, such as T2c1 and U6c1.

The direct analysis of ancient remains from the Canary Islands, using mtDNA by means of PCR techniques, confirmed the presence of North African markers in the indigenous people, including the U6b1a haplogroup, as well as some of the Eurasian lineages observed in the modern population [36]. Admixture analysis based on mtDNA data, using the natives as parental population, determined that 42% of modern Canarian mtDNA lineages have an indigenous origin [36]. Ancient mtDNA results from four of the seven islands found high diversities for Tenerife and La Palma [33,36,37], and the partial and complete fixation of certain haplogroups in La Gomera [38] and El Hierro [39], suggesting that the colonization of the archipelago was a heterogeneous process and that different islands could have had different evolutionary histories.

Although previous ancient DNA (aDNA) studies have been fundamental to understanding the origin and evolution of the Canarian population, most of the ancient mtDNA data produced so far has been obtained using PCR amplification. This classical aDNA technique has provided valuable information, but results have always been hindered by the risk of sample contamination. This is due to the fact that aDNA from warm climates is often extremely degraded and the PCR technique is highly sensitive, thus minute amounts of modern contaminant DNA can be preferentially amplified [40]. Additionally, because the molecules are short and degraded, aDNA analyses based on PCR amplification have tended to isolate small, but informative, regions of the mitochondrial genome, such as the hypervariable region (HVR). This partial information does not allow for refined classification within haplogroups, which is needed to discriminate between close geographical regions. This is especially true within haplogroup H, which comprises ~40% of the ancient Canarian mtDNA lineages. The advent of next-generation-sequencing (NGS) has greatly expanded the capacity of aDNA research. NGS allows damage patterns that are unique to aDNA, such as short fragment size and post-mortem damage, to be detected easily, thus authenticating mtDNA results. NGS also has the advantage of providing complete mtDNA genomes to allow a better geographic assignment, compared to those obtained from partial HVR sequences.

A recent NGS study of the Canarian indigenous people presented the first complete mtDNA genomes and low-coverage full genomes from this population, and, more specifically, from the central islands of Tenerife and Gran Canaria [41]. However, previous aDNA data [36–39] suggested that the indigenous populations from different islands might have experienced different demographic processes. The inclusion of data from all seven islands is therefore of paramount importance to accurately characterizing the archipelago’s indigenous population. Additionally, to fully benefit from the potential of ancient mtDNA data, a more detailed phylogeographic analysis is required.

In order to obtain a comprehensive mtDNA perspective on the origin of the indigenous people of the Canary Islands, we have applied aDNA protocols and NGS to assemble ancient mtDNA genomes from all seven sub-populations. Since human remains from warm regions like the Canary Islands are expected to have low endogenous DNA content, we applied an enrichment technique [42] to improve mtDNA coverage and reduce sequencing costs.

## Methods

### Sample collection

Samples were collected in collaboration with both Canarian universities, La Laguna (Tenerife) and Las Palmas de Gran Canaria (Gran Canaria), as well as the insular museums of Gran Canaria (El Museo Canario), La Palma (Museo Arqueológico Benahorita) and La Gomera (Museo Arqueológico de La Gomera). A total of 25 archaeological sites were selected for this project (Figure 1). Radiocarbon calibrated dates are available for several sites (Figure S1): El Agujero (1030 - 1440 cal AD), La Angostura (1318 - 1394 cal AD), Las Arenas (540 - 650 cal AD), El Capricho (400 - 480 cal AD), Cascajo (1640 - 1700 cal AD), Cuermeja (1270 - 1316 cal AD), La Fortaleza (599 – 633 cal AD), Guayadeque (540 - 737 cal AD), El Hormiguero (1020 - 1160 cal AD), Huriamen (1015 - 1050 cal AD; 1080 - 1150 cal AD), Lomo Galeón (1260 - 1290 cal AD), Montaña Mina (1313 - 1365 cal AD), El Pescante (150 - 350 cal AD), Portillo (1500 - 1580 cal AD), Puente de La Calzada (1265 - 1312 cal AD; 1358 - 1388 cal AD), Punta Azul (1015 - 1155 cal AD) and El Salitre (1060 - 1179 cal AD). For those sites with no available calibrated dates (Antoncojo, Barranco Majona, El Cedro, Cueva del Agua, El Espigón, Huerto de Los Morales, Los Pasitos and Salto del Casimiro), their assignation to the indigenous population was based on general context, the archaeological remains themselves and the presence of specific funerary practices. Sample CAN.005 is a tooth sample that was taken from a private collection of ancient human remains donated to El Museo Canario (Gran Canaria, Spain). Although this sample is not associated with any specific archaeological site, its calibrated radiocarbon date (1265 - 1312 cal AD) is in agreement with a pre-Hispanic origin. It is also worth mentioning that some archaeological sites from Tenerife (Cascajo and Portillo) are from the post-conquest period [43], but they are associated with the so-called “Alzados”, indigenous people that rebelled against the European colonizers and retired to the mountains, leaving all contact with the Europeans behind [44].

### DNA extraction and library preparation

Best-conserved samples were selected for DNA extraction. Although the petrous bone is considered the best source for aDNA [45], we used teeth and small bones (e.g. phalanx) to avoid destroying valuable archaeological material.

Required precautions were taken during the handling of samples, and all experiments that included aDNA were carried out in dedicated, clean lab facilities at the Paleogenomics Lab, University of California Santa Cruz, to avoid contamination. DNA extraction was performed following Dabney et al. [46]. Bone samples were sanded to remove the external surface, and then one bone piece was cut with a Dremel tool and pulverized using a bone mill. The surface of tooth samples was decontaminated using a bleach solution, and then the teeth were cut down the midline and the cementum drilled using a Dremel tool and a metallic bit. Pulverized bone and tooth samples were incubated overnight, using a proteinase K/EDTA solution, and DNA extracted using a silica-based and guanidine method. Ancient DNA was then built into double-stranded libraries, with 7-bp single-index barcoding to allow for multiplexing sequencing, following Meyer and Kircher [47]. Libraries were sequenced for an initial screening on an Illumina NextSeq 500 apparatus for obtaining paired-end shotgun data (~1 M reads per library) with a sequencing read length of 2 x 75 bp.

### Enrichment

After the screening of shotgun libraries, those samples with an endogenous DNA content lower than 10% were enriched using whole-genome in solution capture [42]. Briefly, aDNA libraries were captured in singleplex reactions using human genomic RNA baits, with the aim of increasing endogenous DNA rates and reducing sequencing costs. Although this method is directed at capturing the whole genome, multicopy regions of the mtDNA become particularly enriched. Post-capture libraries were sequenced as indicated before, to obtain at least ~5 M reads per post-capture library.

### HVR analysis

In order to perform population-based analyses, we included in our study previously published [36–39] and unpublished HVR data from the seven islands. Newly reported HVR data from the islands of Gran Canaria (n = 75), Lanzarote (n = 8) and Fuerteventura (n = 10) was obtained following the methodology described by Maca-Meyer et al. [36] and Ordóñez et al. [39]. Briefly, after external decontamination, tooth samples were extracted by means of a GuSCN-silica based protocol. MtDNA quantification was performed on a 7500 Real Time PCR system (Applied Biosystem, Foster City, CA, USA), using a human-specific mtDNA fluorescent probe [48], and ~3,000 copies were submitted to PCR with the aim of reducing the effects of DNA damage. The mtDNA HVRI (from positions 16,000 to 16,400) was amplified using seven overlapping fragments, with sizes ranging from 82 to 124 bp, to improve the amplification of endogenous DNA. All the sequencing reactions were prepared with the BigDye v3.1 Terminator Cycle Sequencing kit (Applied Biosystems) and run on an ABI PRISM 3130xl Genetic Analyzer (Applied Biosystems). Standard contamination prevention and monitoring were conducted as described earlier [39].

### Modern mtDNA genomes

We included in this study several current Canary Islands mtDNA genomes, analyzed using both whole-genome and Sanger sequencing. Complete genomes were obtained in Instituto Tecnológico y de Energías Renovables (ITER) by whole-genome sequencing from a set of 18 unrelated Canarians. Briefly, DNA samples were processed with a Nextera DNA Prep kit, with dual indexes following the manufacturer’s recommendations (Illumina Inc., San Diego, CA). Library sizes were checked on a TapeStation 4200 (Agilent Technologies, Santa Clara, CA) and their concentration determined by the Qubit dsDNA HS Assay (Thermo Fisher, Waltham, MA). Samples were sequenced to a depth of 30X on a HiSeq 4000 instrument (Illumina) with paired-end 150-base reads. Sanger sequencing mtDNA genomes were obtained at University of La Laguna following previously published methodologies [49], for samples classified as T2c1 (determined by HVRI analysis). These samples were selected because of their potential to define new sub-lineages within T2c1. Institutional review board approval for the analysis of human subjects was obtained from Stanford University.

### Data analysis

#### Mapping and filtering of ancient mtDNA reads

Shotgun sequencing reads were trimmed and adapters removed using AdapterRemoval version 1.5.4 [50]. Specifically, the paired-end reads were merged, and low-quality bases (BASEQ < 20) and short reads (< 30 bp) removed. Merged trimmed reads were then mapped to the human reference genome (hg19) using BWA version 0.7.12 [51], while unmerged reads were discarded. Unmapped, low-quality (MAPQ<30) and duplicate reads were removed using SAMtools version 0.1.19 [52]. The percentage of endogenous DNA was calculated by dividing the number of reads remaining after filtering by the total number of trimmed reads.

#### Authentication

Damage patterns were assessed using MapDamage v2.0 [53]. Insert size of libraries was obtained with SAMtools mpileup, and plotted using R software v.3.2.0 [54]. Contamination rates of libraries were calculated using contamMix v.1.0-10 [55] and Schmutzi [56].

#### Analysis of complete mtDNA genomes

MtDNA reads were directly mapped to the revised Cambridge Reference Sequence (rCRS) [57] and filtered as described before. MapDamage was used to rescale the quality of bases likely affected by post-mortem damage. Indel Realigner from the GATK pipeline version 2.5.2 was also used for improving alignment quality around indels [58]. MtDNA consensus sequences were obtained using SAMtools and BCFtools version 0.1.19 [52]. A list of variants was then obtained using SAMtools mpileup, with a minimum depth of 5. Haplogroups were determined with HaploGrep version 2.0 [59], using PhyloTree build 17 version (http://www.phylotree.org) [60]. MtDNA haplotypes were manually curated by visual inspection, using Tablet v.1.17.08.17 [61]. Modern DNA sequencing data was analyzed following the same protocol used for ancient samples, except for the MapDamage rescaling step. After retrieving all available mtDNA genomes belonging to the haplogroups of interest from NCBI (http://www.ncbi.nlm.nih.gov), phylogenetic trees were built using median-joining networks [62]. Indels around nucleotides 309, 522, 573 and 16193, and hotspot mutations (e.g. 16519) were excluded from phylogenetic analysis. For estimating coalescence ages for specific clades, we used the ρ statistic [63]. We established a mutation rate for the complete mtDNA sequence of one substitution in every 3,624 years, correcting for purifying selection as in Soares et al. [64]. Accompanying standard errors were calculated as per Saillard et al. [65]. For highly frequent haplogroups, such as H1cf and T2cd3, we only kept one sample per site, to avoid relatedness interfering with coalescence age estimations.

#### Analysis of HVRI data

Newly reported HVR sequences were analyzed using BioEdit software v.7.0.9.0 [66], and haplotypes were obtained by means of HaploSearch software [67] and further confirmed by manually inspecting the electropherograms. Haplogroup nomenclature was assessed following the most updated mtDNA phylotree (Build 17) [60].

Genome-wide data was combined with previous HVRI sequencing data to perform population-based analysis. Published samples used for comparisons are detailed in Table S1. As we do not know if samples in the same burial can be related, when several samples with the same haplogroup were observed from the same archaeological site, only one was included in the analysis. Two-tailed Fisher’s exact test was used to assess differences in mtDNA haplogroup frequencies between eastern and western islands. Gene diversity was calculated according to Nei [68]. Distances between populations were estimated using haplogroup frequency-based linearized FST [69] as in Arlequin v.3.5 [70]. Multidimensional Scaling (MDS) was performed using R software and the “smacof” package [71]. Admixture estimates were calculated with the WLSAdmix program [72], which was kindly provided by Dr Jeffrey Long.

## Results and Discussion

The average endogenous DNA content for the Canarian indigenous samples is 7.92%, a relatively high value considering the warm and humid environmental conditions of the archipelago (Table S2). However, endogenous DNA values varied within and between archaeological sites, ranging between 0.02% and 39.0% (IQR= 0.67% - 11.5%). All samples meet the standard aDNA authentication criteria, including observation of DNA fragmentation and damage patterns at both ends of molecules, and low modern DNA contamination rates (Figure 2). Those contamination rates calculated with contamMix are larger than those produced with Schmutzi. One possible reason is that contamMix estimations are more sensitive to low coverage values (Table S2). For example, sample CAN.033, with a 7.9X mtDNA coverage, has a contamination rate of 10.2% based on contamMix and 1.0% on schmutzi. Schmutzi has been reported to be able to obtain accurate contamination rates for coverage down to ~5X [56]. However, in other cases, variable contamination estimations do not seem to be related to low coverage, and other factors may be interfering.

**Figure 2.**
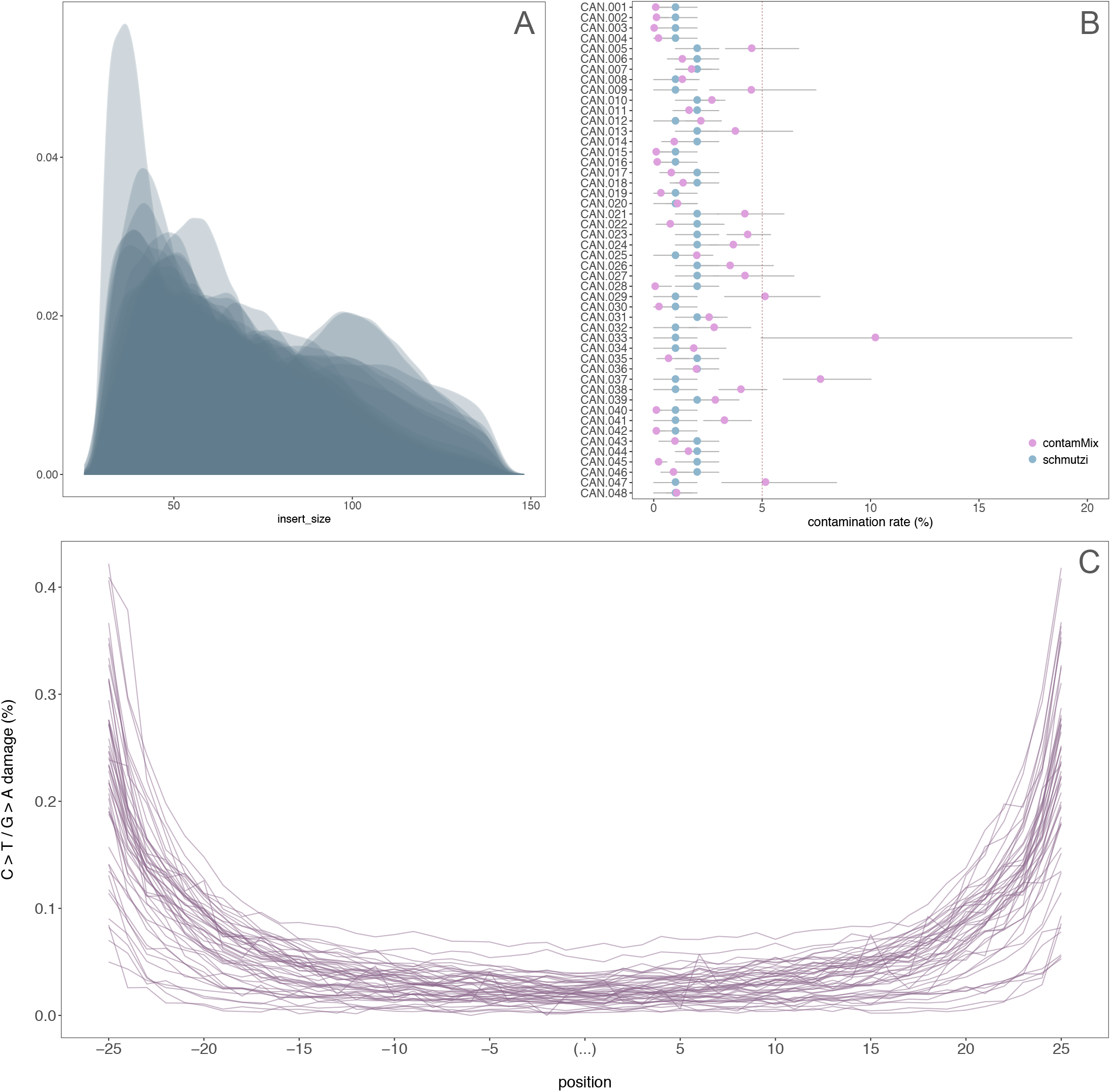
DNA authentication results for all the samples included in this study. A) Insert size density plot. B) Contamination rates estimated using contamMix and schmutzi. C) Damage patterns.

After capture, we obtained complete mtDNA genomes from 48 ancient human remains sampled in 25 different archaeological sites (Table S2). Our sample set covers the entire archipelago and a time span of 1,200 years (Figure S1). The average mtDNA depth is ~140X, with a minimum value of 8X (Table S2). Observed haplogroups agree with previous studies [33,36–39,41], indicating the presence of North African (U6), Eurasian (H, J2, T2 and X) and sub-Saharan African lineages (L1 and L3) in the Canarian indigenous population (Figure S2). As delineated before [36], the majority of haplogroups observed are of Eurasian origin, most with a Mediterranean distribution. This result is expected, as recent aDNA data from North Africa has indicated the presence of Neolithic European lineages as early as the Late Neolithic period (~5,000 BP) [73].

We also obtained complete mtDNA genomes from a set of 18 modern Canarians (Figure S3). More than 50% of the samples belong to haplogroup H, with a higher diversity of sub-haplogroups than the one observed in the indigenous population. In addition to H1cf and H1e1a, we observe other H1 sub-lineages and other branches, such as H6a1, H3c2 or H43, which are most likely of European origin. Other haplogroups present in the indigenous people are also observed in the modern population, including J2a2d, U6b1a and X3a. In line with previous analyses [27,31], a sub-Saharan African (L3d1b3a) [74] and an Amerindian lineage (A2) [75] are observed in the current population of the Canary Islands. Assuming that our set of 48 ancient genomes is representative of the native population, we performed a rough admixture estimate of 27.8% of maternal lineages in members of the present-day population possessing indigenous origins, while 61.1% would be of European ascription (Figure S3).

### Population-based analysis

In order to compare our samples to previously published data, we combined the newly generated mtDNA genomes with HVRI data from the Canarian indigenous population (Table S1) [33,36–39]. Given that sample sizes for Lanzarote and Fuerteventura are small and their indigenous populations are considered to be similar based on archaeological data [76], these data sets were pooled together. It is worth mentioning that those samples for which mitochondrial data were generated, using both classical techniques and NGS sequencing, produced identical HVRI haplotypes, proving our PCR-based approach generates authentic results.

As previously observed, the indigenous populations of the Canary Islands in the past were not homogenous (Table 1; Figure 3). The islands of La Palma and Tenerife show a relatively diverse mtDNA composition (>70%) [33,36,37], while the others show signs of genetic drift and/or diversity reduction events, such as a bottleneck or a founder effect. In La Gomera, mtDNA diversity was 54.2%, due to the high frequency of haplogroup U6b1a [38], while in El Hierro, this value was 2.9%, with the almost complete fixation of H1cf haplogroup in the Punta Azul site [39]. With new data on the indigenous population of Gran Canaria, Lanzarote and Fuerteventura (Table S3), we show that Gran Canaria had high mtDNA diversity, similar to Tenerife and La Palma, while Lanzarote and Fuerteventura had low diversity (51.1%) because of the high frequency of H*(xH1cf, H4a1a) lineages. These findings emphasize that results obtained from the larger islands of Tenerife and Gran Canaria should not be extrapolated to the entire archipelago. Estimations of population sizes during pre-colonial times based on archaeological evidence agree with mtDNA results. Populations in Gran Canaria, Tenerife and, to a lesser degree, La Palma, were large and able to sustain relatively high diversity, while Lanzarote, Fuerteventura and El Hierro were almost depopulated at the time of the conquest [77]. In the case of La Gomera, the population size was also reported to be small [78].

**Table 1.** MtDNA haplogroup absolute frequencies for the indigenous population of the Canary Islands. Haplogroup frequencies and diversity were calculated using HVRI sequence data from this study and previously published data. 1: This study; 2: Ordóñez et al. 2017; 3: Fregel et al. 2009; 4: Maca-Meyer et al. 2004; 5: Fregel et al. 2014; 6: Rodríguez-Varela et al. 2017.

| Haplogroup | El Hierro<sup>1,2</sup> | La Palma<sup>1,3</sup> | Tenerife<sup>1,4,6</sup> | La Gomera<sup>1,5</sup> | Gran Canaria<sup>1,6</sup> | Lanzarote & Fuerteventura<sup>1</sup> | Total |
| --- | --- | --- | --- | --- | --- | --- | --- |
| H* | 12 | 10 | 15 | 2 | 33 | 13 | 85 |
| H1cf | 57 | 8 | 6 | 2 | 1 | - | 74 |
| H4a1e | - | - | - | - | 3 | 1 | 4 |
| HV0 | - | - | 1 | - | - | - | 1 |
| J | - | 3 | 4 | 5 | 2 | - | 14 |
| K | - | 1 | 1 | 1 | 2 | - | 5 |
| L1/L2 | - | 2 | 2 | 1 | 1 | - | 6 |
| L3 | - | - | 2 | 4 | 1 | - | 7 |
| L3b1a12 | - | - | - | - | 5 | - | 5 |
| HV0 | - | - | - | - | 1 | - | 1 |
| Other T | - | 1 | - | - | 3 | - | 4 |
| T2c1 | - | 3 | 12 | - | 15 | 2 | 32 |
| U5 | - | - | - | - | 3 | 1 | 4 |
| U6a | - | - | 2 | - | 6 | 2 | 10 |
| U6b | - | 2 | 8 | 38 | 4 | - | 52 |
| U6c | - | - | - | - | 5 | 1 | 6 |
| U7 | 1 | - | - | - | - | - | 1 |
| W1e1 | - | 1 | - | - | - | - | 1 |
| X3a | - | 4 | - | 4 | 1 | - | 9 |
| Sample size | 70 | 35 | 53 | 57 | 86 | 20 | 321 |
| Haplogroup diversity | 2.86% ± 2.76% | 72.10% ± 7.63% | 77.43% ± 4.02% | 54.20% ± 7.50% | 77.10% ± 3.78% | 51.05% ± 12.84% | 69.69% ± 2.37% |

**Figure 3.**
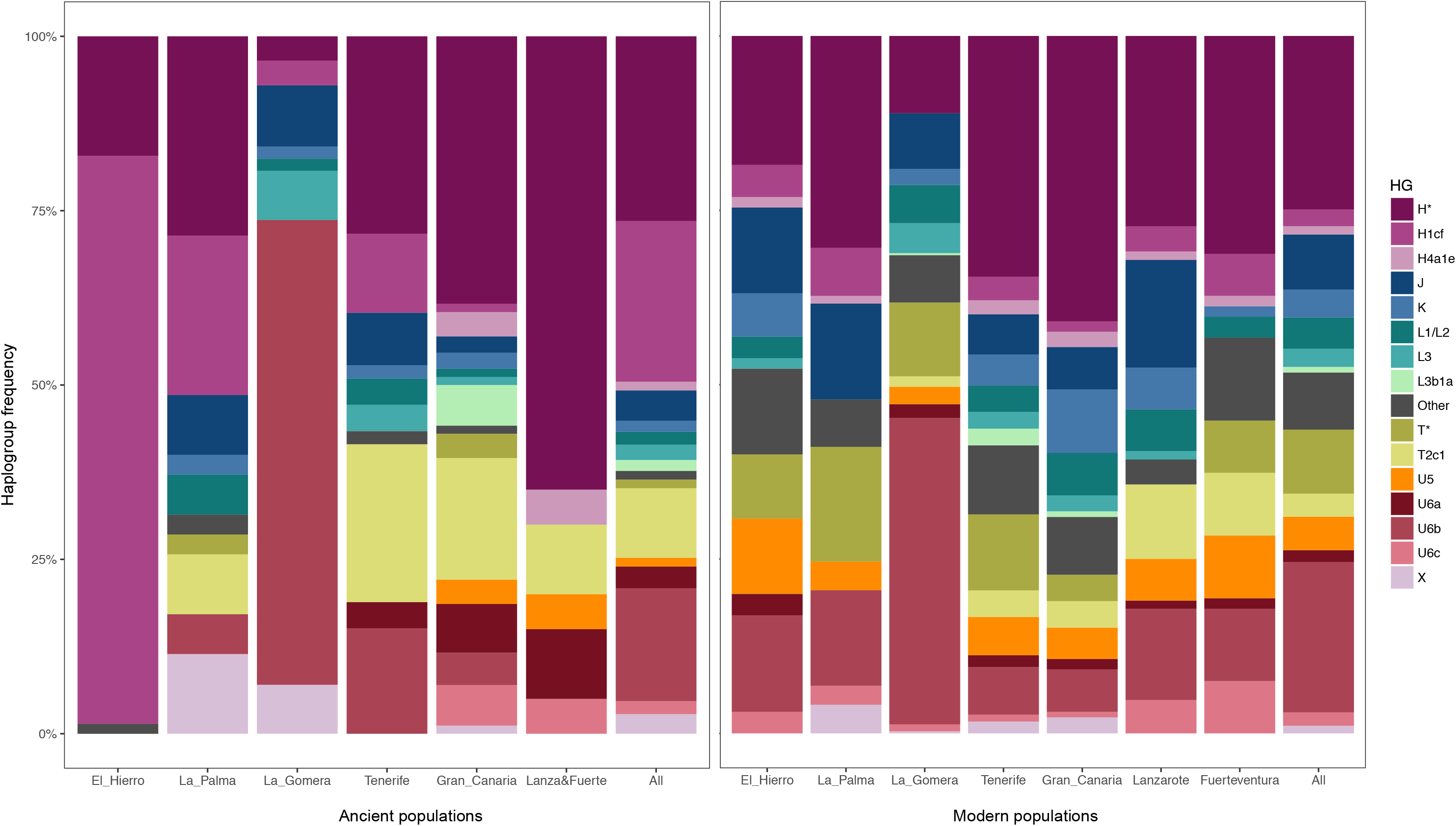
MtDNA haplogroup frequencies for ancient and current populations of the Canary Islands.

By directly comparing the mtDNA types found in the indigenous population of each island, we observe that H4a1e, L3b1a, U5 and U6c haplogroups are present only in the eastern islands (Gran Canaria, Lanzarote and Fuerteventura). Differences between eastern and western islands were shown to be significant for the four haplogroups, when all the ancient samples were considered: H4a1 (*p*=0.0127), L3b1a (*p*=0.0037), U5 (*p*=0.0114) and U6c (*p*=0.0012). Though also present in the western islands, haplogroups T2c1 (*p*=0.0164) and U6a (*p*=0.0028) appeared more frequently in the eastern islands. However, these results can be artifacts caused by the high frequency of H1cf in El Hierro and U6b1a in La Gomera. After removing these two populations from the western group, only differences in the distribution of U6c remained significant (*p*=0.0328).

In contrast with the heterogeneity we observe in pre-Hispanic times, mtDNA haplogroup frequencies in modern populations of the Canary Islands are homogenous (Figure 3), with the sole exception of La Gomera [27]. The high frequency of haplogroup U6b1a observed in the indigenous population of La Gomera is also detected in its present-day population [38]. However, the same pattern is not observed for El Hierro. In pre-colonial times, H1cf was almost fixated in El Hierro [39], while the frequency of this haplogroup today is 4.6%, not significantly different from the average 2.4% observed in the entire archipelago (*p*=0.2364).

In order to determine the admixture pattern at an insular level, we compared modern Canarian samples with their principal parental populations: indigenous people, Iberians, and sub-Saharan Africans (Table S1). Global admixture estimations using the new mtDNA dataset (Table 2) confirm previous results on the survival of native lineages in the modern population (55.9%). However, we observed that results within islands are variable. When the miscellaneous ancient sample is used as one of the parental populations, indigenous contribution to the modern population ranges from 30.8% in Gran Canaria to 71.4% in La Gomera. However, this approach is not correct, as we know that the indigenous population of the archipelago was heterogeneous and mtDNA frequencies were variable. With our new data, we were able to estimate admixture, using aDNA sampled directly from each island. Indigenous mtDNA contribution estimates are lower when a direct comparison is performed, with values ranging from 0% in El Hierro to 55.5% in La Gomera (Table 2). The extreme result observed in El Hierro is evidently due to the marked difference between the ancient and current people. It is interesting that, when the miscellaneous sample is used, the indigenous contribution increases to 36.2%. This result is reasonable, given that the present-day sample from El Hierro is not significantly different from other islands. This can be explained if we consider that El Hierro was almost depopulated at the time of the European conquest [79]. In fact, it was recounted in the chronicles that the indigenous population of El Hierro was decimated due to razzias (raids for the purpose of capturing slaves) at the time of the Spanish conquest, and was later repopulated with indigenous populations from other islands and European colonizers [80,81].

**Table 2.** Admixture results based on mtDNA haplogroup frequencies. Admixture results for the modern population of the Canary Islands using the three main parental populations: Iberian Peninsula (IBP), sub-Saharan Africa (SSA) and the Canarian indigenous population (CIP). Admixture calculations were performed using two approximations: A) we used the whole ancient dataset (combining the ancient samples from all the seven islands) as CIP for calculating admixture estimates for all islands; B) we used each indigenous sample to calculate the admixture of its respective island (e.g. to calculate admixture in the modern population of Gran Canaria we exclusively used the ancient samples from Gran Canaria as CIP). Results are shown for: the whole Canary Islands population (CAN) and the seven individual islands (FUE=Fuerteventura; GCA=Gran Canaria; GOM=La Gomera; HIE=El Hierro; LAN=Lanzarote; PAL=La Palma; TFE=Tenerife).

| **A: Whole indigenous sample** |  |  |  |  |  |  |  |  |  |
| --- | --- | --- | --- | --- | --- | --- | --- | --- | --- |
| **Component** | **IBP** |  |  | **SSA** |  |  | **CIP** |  |  |
| **FUE** | 0.4108 | ± | 0.0071 | 0.0199 | ± | 0.0015 | 0.5692 | ± | 0.0070 |
| **GCA** | 0.6486 | ± | 0.0039 | 0.0438 | ± | 0.0012 | 0.3076 | ± | 0.0038 |
| **GOM** | 0.2170 | ± | 0.0181 | 0.0691 | ± | 0.0075 | 0.7139 | ± | 0.0186 |
| **HIE** | 0.6379 | ± | 0.0122 | 0.0000 | ± | 0.0000 | 0.3621 | ± | 0.0122 |
| **LAN** | 0.3303 | ± | 0.0084 | 0.0448 | ± | 0.0026 | 0.6248 | ± | 0.0084 |
| **PAL** | 0.5599 | ± | 0.0106 | 0.0000 | ± | 0.0001 | 0.4401 | ± | 0.0107 |
| **TFE** | 0.5989 | ± | 0.0041 | 0.0452 | ± | 0.0012 | 0.3559 | ± | 0.0040 |
| **CAN** | 0.3978 | ± | 0.0100 | 0.0432 | ± | 0.0029 | 0.5589 | ± | 0.0100 |
| **B: Indigenous sample from each island** |  |  |  |  |  |  |  |  |  |
| **Component** | **IBP** |  |  | **SSA** |  |  | **CIP** |  |  |
| **FUE** | 0.6458 | ± | 0.0095 | 0.0203 | ± | 0.0020 | 0.3339 | ± | 0.0093 |
| **GCA** | 0.6944 | ± | 0.0040 | 0.0593 | ± | 0.0015 | 0.2463 | ± | 0.0038 |
| **GOM** | 0.3768 | ± | 0.0049 | 0.0682 | ± | 0.0024 | 0.5550 | ± | 0.0049 |
| **HIE** | 1.0000 | ± | 0.0087 | 0.0000 | ± | 0.0000 | 0.0000 | ± | 0.0087 |
| **LAN** | 0.7202 | ± | 0.0116 | 0.0239 | ± | 0.0027 | 0.2559 | ± | 0.0113 |
| **PAL** | 0.5896 | ± | 0.0134 | 0.0000 | ± | 0.0116 | 0.4104 | ± | 0.0118 |
| **TFE** | 0.7306 | ± | 0.0030 | 0.0495 | ± | 0.0012 | 0.2199 | ± | 0.0029 |

To determine if a more specific origin for the Canarian indigenous population could be ascertained, the ancient mtDNA sample was combined with a reference modern DNA database containing samples from the Canary Islands, Europe, North Africa, Sub-Saharan Africa and the Near East (Table S1). In the MDS analysis (Figure 4), the indigenous sample from El Hierro and the indigenous and modern samples from La Gomera act as outliers, due to the high frequency of H1cf and U6b1a, respectively. When the two outliers were removed and all the remaining ancient samples were pooled together, the first dimension differentiates sub-Saharan populations from Eurasian populations, including North Africa and the Canary Islands. The second dimension places Canarian and European/Near Eastern populations on both ends, with North Africans in an intermediate position. The closest North African sample to the Canarian indigenous population in the second dimension is West Sahara. However, the ancient sample is differentiated from all current North African populations and placed closer to modern Canarians. This is due to the fact that haplogroups occurring frequently in the Canarian indigenous and current samples (e.g. U6b1a) are not present or appear in low frequencies within the reference populations. This result concurs with later demographic processes reshaping the mtDNA landscape of North Africa, and/or founder effects and isolation in the Canary Islands. It is interesting that, compared to the other islands, the modern populations of Tenerife and Gran Canaria are closer to the European populations. This result is expected, because they each have capital cities of the two Canarian provinces and, thus, have received substantial historical migration from the mainland.

**Figure 4.**
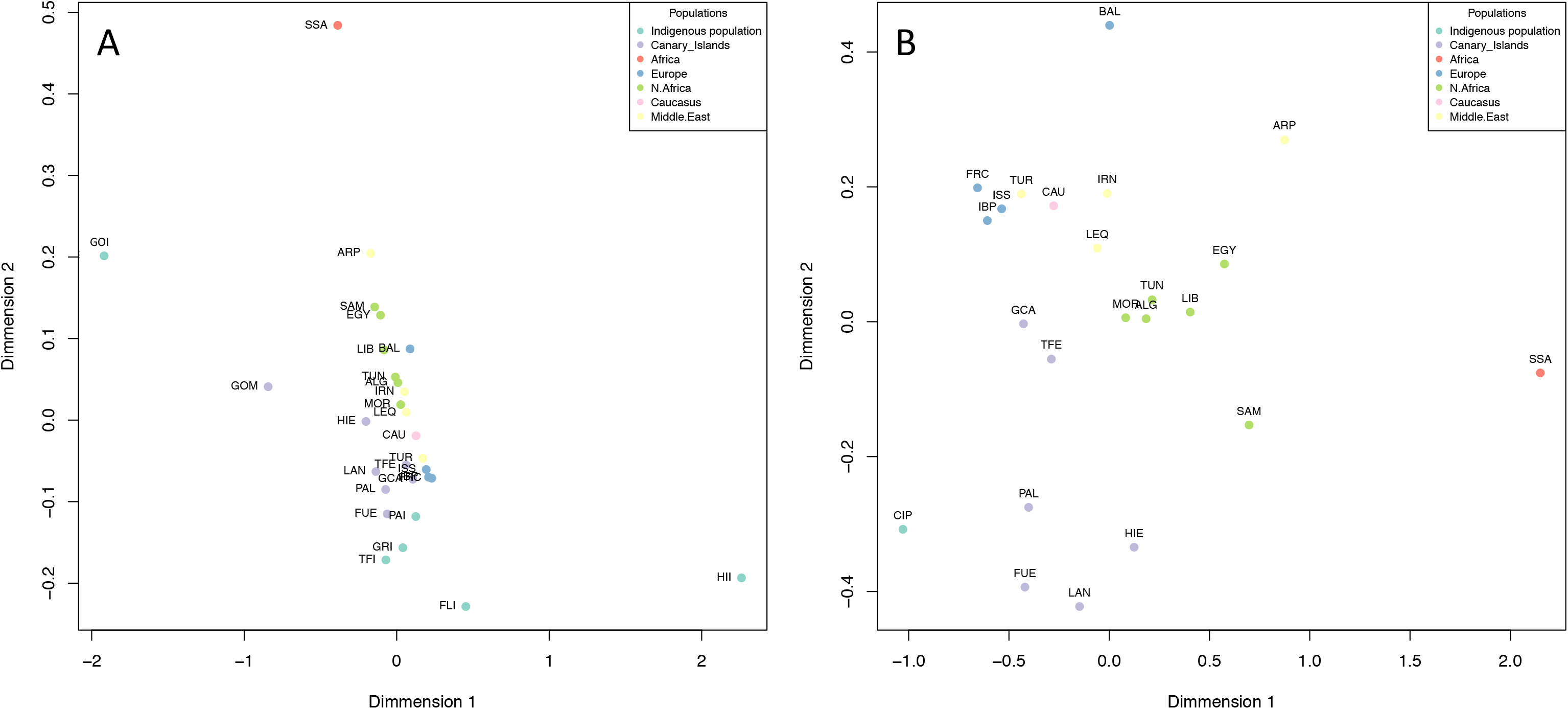
MDS plot based on haplogroup frequency distances. A) MDS analysis comparing the individual ancient populations (FUI=Fuerteventura; GCI=Gran Canaria; GOI=La Gomera; HII=El Hierro; LAI=Lanzarote; PAI=La Palma; TFI=Tenerife), with modern Canarian (codes as in Table 2), Caucasus (CAU), North African (codes as in Table S1), Sub-Saharan African (SSA), European (codes as in Table S1) and Near Eastern populations (codes as in Table S2). B) MDS analysis as in A, but removing outliers (HIE, HII and GOM) and pooling all the remaining indigenous samples together (CIP).

### Phylogeographic analysis of mitogenomes

The HVRI has been proven to be of limited value in providing a clear picture of the origin of the indigenous people of the Canary Islands. In order to conduct a better assignment of the geographic origin of the maternal Canarian indigenous lineages, we performed detailed phylogeographic analysis of all the lineages observed in the aDNA dataset (Figure S2), including those from Rodriguez-Varela et al. [41]. For detailed information on our phylogenetic analysis, see Supplementary Text.

We observe five different H sub-lineages in the indigenous people of the Canary Islands: H1cf, H1e1a9, H2, H3 and H4a1e. H1cf (Figure S4) seems to be restricted to both the Canary Islands and Central North Africa, and shows a coalescence age (~3,400 years ago) that is in agreement with a continental origin before the colonization of the islands (Figure S5). Newly defined haplogroups H1e1a9 (Figure S6) and H4a1e (Figure S7) are both restricted to the Canary Islands, with a distribution similar to that observed for U6b1a. However, in this case, H1e1a9 and H4a1e coalescence ages overlap with the human occupation period (Figure S5) and are compatible with an origin in the islands. The presence of lineages derived of H1e1a and H4a1 in both European Neolithic and the Canary Islands indigenous samples corresponds with Eurasian prehistoric intrusions in North Africa (Fregel et al. 2018). Two samples were classified within basal H2 and H3 haplogroups, preventing further phylogenetic analysis.

Two sublineages of haplogroup J are observed in the indigenous population of the Canary Islands: J1c3 and the newly defined J2a2d1a1. J1c3 is present in Europe, North Africa and the Near East, and more interestingly, in ancient Neolithic samples from Spain and Sardinia (Figure S8). Although J2a2d1a* has been spotted in Central North Africa, subhaplogroup J2a2d1a1 is exclusive to the Canary Islands and Brazil, the latter representing an area with known historical migrations from the islands (Figure S9). Accordingly, this new autochthonous Canarian lineage has a coalescence age that overlaps with the indigenous occupation of the islands (Figure S5).

Phylogenetic analysis of the Canarian T2 sequences places them within T2b and T2c1d, two haplogroups thoroughly observed in Neolithic and Bronze Age sites from Europe. The inclusion of ancient and modern Canarian samples allows us to define four new T2c1d subhaplogroups (Figure S10). T2c1d3 haplogroup is present in both Tunisia and the Canary Islands. T2c1d1c1 and its two subclades (T2c1d1c1a and T2c1d1c1b) are present in both North Africa and the current population of the eastern Canary Islands. This distribution could be explained by an asymmetrical migration pattern, or, given its absence in the indigenous people, by a higher impact of Moorish slave trade in the eastern islands (Supplementary Text).

We identify several indigenous samples within macrohaplogroup L, belonging to L1b1a and the newly defined L3b1a12. Although Later Stone Age [82], and Early and Late Neolithic [73] samples from North Africa did not show any mtDNA lineage of sub-Saharan origin, our results imply the presence of L1b and L3b1a in North Africa at the time of the colonization of the Canary Islands. Regarding L3b1a12 (Figure S11), this lineage can also be considered autochthonous of the Canary Islands, with a coalescence age posterior to the proposed colonization date (Figure S5). Interestingly, this lineage was only present in the eastern islands in ancient times, but has a wider distribution at the present time, suggesting extensive movement of native people after the conquest.

Canarian indigenous sequences belonging to X haplogroup are classified within the X3a clade (Figure S12). This lineage is present both in Europe, the Near East and northeast Africa, as well as in the ancient and current populations of the Canary Islands.

Finally, several U6 sublineages are observed in the indigenous population of the Canary Islands, including U6a1a1 (Figure S13), U6a7a1 (Figure S14), U6b1a (Figure S16) and U6c1 (Figure S16). U6a1a1, U6a7a1 and U6c1 are present in the Maghreb, southern Europe and the Canary Islands, and are most probably related to prehistoric Mediterranean expansions (Figure S13, Figure S14 and Figure S16). As reported before, the Canarian autochthonous U6b1a is also present in regions with recent Canarian migration, including mainland Spain and Cuba (Figure S15). Given its coalescence age and the oldest calibrated radiocarbon dates from human remains from the Canary Islands (Figure S5), U6b1a most probably originated in North Africa and later migrated to the Canaries. However, to date, this lineage has not been observed in the continent, indicating the migrations occurred after the colonization of the Canary Islands reshaped the North African mtDNA landscape.

## Discussion

Our mtDNA results on the indigenous people of the Canary Islands shed light on the prehistory of North Africa. Our data are in agreement with recent aDNA data from Morocco [73] and further evidence of a complex pattern of Mediterranean migrations in North Africa. Archaeological records in the Maghreb support this result, and also suggest further European intrusions during the Chalcolithic and Bronze Age eras [83,84]. Additionally, Phoenicians, Carthaginians and Romans arrived in the North African region in historical times [85–88]. The presence of haplogroups of Mediterranean distribution in the indigenous people of the Canaries demonstrates the impact of these prehistoric and historical migrations in the Berbers and that they were already an admixed population at the time of the indigenous colonization of the islands [89].

In our phylogeographic analysis of complete mtDNA sequences from the Canarian indigenous population, we found lineages that are only observed in Central North Africa and the Canary Islands (H1cf, J2a2d and T2c1d3), while others have a wider distribution including both West and Central North Africa, and, in some cases, Europe and the Near East (U6a1a1, U6a7a1, U6b, X3a, U6c1). These results point to a complex scenario, where different migration waves from a dynamic and evolving North African population reached the islands over time. Every island experienced their own evolutionary path, determined by the environmental conditions and limitations of insularity. Those islands with the capability of sustaining large populations retained variability, while others with more restricted means (La Gomera and probably El Hierro) had to develop cultural practices to avoid inbreeding, like mandatory exogamic practices [78,90].

Although the North African Berber origin is the most widely accepted hypothesis, other lines of research have proposed that certain funerary practices and religious beliefs observed in the indigenous population of the Canary Islands could be linked to Punic-Phoenician influence [91], thus proposing the colonization of the Canary Islands as the result of Phoenicians expanding their control to the Atlantic Ocean. Based on the limits of the territorial occupation of the Atlantic West Africa by Phoenicians, Carthaginians and Romans, most researchers consider it unlikely that there was a political occupation or economic exploitation of the archipelago [92–94]. However, the islands were not unknown to Mediterranean cultures, and Romans possessed the seafaring skills needed to travel to the islands [22]. Some authors think Phoenicians also had the navigational technology required to reach the Canary Islands [95,96], although this idea has been challenged [97]. The first Phoenician aDNA sample published was a complete mtDNA sequence of a child from Carthage dated to the 6^th^ century BC [98]. This Carthaginian sample was classified within U5b2c1 haplogroup. This result is interesting, given that U5 was more frequent in the indigenous population of the eastern islands, including the island of Lanzarote, where a Punic-Phoenician influence has been proposed. As U5 haplogroup was not uncommon in Neolithic European samples, and its presence in North Africa might be due to prehistoric migrations, an alterative explanation would be that haplogroup U5 was incorporated into the Berber mtDNA pool before the Carthaginians were established in Tunisia. Recently, Matisoo-Smith et al. [99] published thirteen complete mitogenomes from Punic-Phoenician samples from Lebanon and Sardinia. The only haplogroups in common with the indigenous population of the Canary Islands are H3 and H1e1a, although, in this case, the Phoenician H1e1a sample is classified within the sub-lineage H1e1a10. The lack of overlap between the mtDNA composition of Phoenicians and the Canarian indigenous people disagrees with either a Punic-Phoenician origin for the ancient islanders or sustained contact between the two populations.

Previous genetic analyses of the modern Canarian population detected an asymmetrical distribution of maternal and paternal lineages in the archipelago [31,35]. Our aDNA results confirm the existence of asymmetrical distribution of mtDNA haplogroups in pre-colonial times, with the presence of haplogroups H1e1a9, H4a1e, L3b1a12 and U6c1 only in the eastern islands. However, it is worth mentioning that La Palma, the island with the most anthropological evidence of two migrations waves, does not show any of these lineages. If we consider the presence of H1e1a9, H4a1e, L3b1a12 and U6c1 haplogroups to be the result of further population movements from North Africa to the eastern islands, we could approximate the date based on radiocarbon dates of the sites where the sample was taken. Most sites where these lineages have been observed have radiocarbon dates placed around the 13^th^ century, and all except one are from after the 10^th^ century. The only site with an older date is Guayadeque; however, we have to take into account that this is a large site, with evidence of human occupation extending until the 14^th^ centuries AD [100], and the dating was not performed directly on the analyzed sample.

Archaeological data has evidenced significant changes in the productive strategies of some islands around the 11^th^ - 12^th^ centuries [12,76,101–103]. In fact, recent data indicates probable population growth in Gran Canaria at that time, suggesting the appearance of new settlements associated with an exploitation model that intensified the use of marine resources, the increase in the size of settlements linked to agricultural nuclei, and changes in the production of some craftsmanships [12,104,105]. These changes have been interpreted as part of an endogenous process, as it has been determined that this population growth involved neither significant changes in the structure of human settlements or burials, nor introduced differences in land management or the types of domestic species that were exploited. However, it is also possible to explain those changes as the result of the arrival of new migrants to the island of Gran Canaria. Although it is still under study, there is evidence for transformations in the configuration of some settlements in Lanzarote, between the 8^th^ and 13^th^ centuries [106]. Again, these modifications could be reflecting changes in the conception of domestic space due to an endogenous process, or associated with the arrival of new colonizers. Archaeological information from Fuerteventura is not abundant enough to determine population size changes that could be related to the arrival of new migrants. Nevertheless, it is clear from the archaeological record that Fuerteventura and Lanzarote maintained frequent contact and shared both cultural and economic elements [76,107]. Future paleogenomic efforts to obtain high-coverage genomes from all seven islands, in combination with proper archaeological contextualization of the genetic data and detailed radiocarbon dating, will be essential for improving our knowledge of the origins and evolution of the indigenous population of the Canary Islands.

## Supporting information

## DATA AVAILABILITY

Mitochondrial DNA sequence data are available through the European Nucleotide Archive (PRJEB29569). Consensus mtDNA sequences are available at the National Center of Biotechnology Information (Accession Numbers MK139577 - MK139649). Requests for additional materials should be addressed to R.F.

## ACKNOWLEDGMENTS

C.D.B. and R.F. were funded by a grant from the National Science Foundation (1201234) and the Chan Zuckerberg Biohub Investigator Award; R.F. was funded by a Fundación Canaria Dr. Manuel Morales Fellowship and by a grant from Dirección General de Patrimonio Cultural del Gobierno de Canarias (MITOCAN); B.S. was funded by a grant from the Gordon and Betty Moore Foundation (GBMF-3804); R.G.M., J.M.L.S and C.F. were funded by grants from the Spanish Ministry of Science and Innovation (RTC-2017-6471-1) and Área Tenerife 2030 from Cabildo de Tenerife (CGIEU0000219140), and from the agreement OA17/008 with Instituto Tecnológico y de Energías Renovables (ITER). Finally, C.F. wants to acknowledge technical assistance from Ana Díaz-de-Usera.

## TITLES AND LEGENDS TO SUPPLEMENTARY MATERIAL

**Table S1 – Populations used for comparisons in this study.**

**Table S2 – Summary of mtDNA results for all aDNA samples.**

**Table S3 – HVRI data used for this study, including new results on the islands of Gran Canaria, Lanzarote and Fuerteventura.**

**Figure S1 – Combined calibrated radiocarbon per archaeological site (A) and per mtDNA lineage (B).**

**Figure S2 –** Phylogenetic tree of complete ancient Canarian mtDNA sequences.

Number along links refers to nucleotide changes, whereas “@”, “d” and “i” indicates back mutations, deletions and insertions, respectively. Recurrent mutations, such as 309iC, 315iC and 16519, have not been taken into account.

**Figure S3 – Phylogenetic tree of complete modern Canary Islands sequences.** The most probable geographic origin of the sequences is indicated.

**Figure S4 – Phylogenetic tree of complete haplogroup H1cf sequences.** GenBank accessions and geographic origin are indicated for each complete sequence taken from the bibliography.

**Figure S5 – Coalescence ages for mtDNA haplogroups observed in the indigenous population of the Canary Islands.** All the coalescence ages have been calculated in this study, except for H2a, H3 and T2b, whose ages have been obtained from previous results (Behar et al. 2008).

**Figure S6 – Phylogenetic tree of complete haplogroup H1e1a sequences.** GenBank accessions and geographic origin are indicated for each complete sequence taken from the bibliography. Sub-haplogroups in dark grey and white fonts indicate newly defined branches.

**Figure S7 – Phylogenetic tree of complete haplogroup H4a1 sequences.** GenBank accessions and geographic origin are indicated for each complete sequence taken from the bibliography. Sub-haplogroups in dark grey and white fonts indicate newly defined branches.

**Figure S8 – Phylogenetic tree of complete haplogroup J1c3 sequences.** GenBank accessions and geographic origin are indicated for each complete sequence taken from the bibliography.

**Figure S9 – Phylogenetic tree of complete haplogroup J2a2d sequences.** GenBank accessions and geographic origin are indicated for each complete sequence taken from the bibliography. Sub-haplogroups in dark grey and white fonts indicate newly defined branches.

**Figure S10 – Phylogenetic tree of complete haplogroup T2c1d sequences.** GenBank accessions and geographic origin are indicated for each complete sequence taken from the bibliography. Sub-haplogroups in dark grey and white fonts indicate newly defined branches.

**Figure S11 – Phylogenetic tree of complete haplogroup L3b1a sequences.** GenBank accessions and geographic origin are indicated for each complete sequence taken from the bibliography. Sub-haplogroups in dark grey and white fonts indicate newly defined branches.

**Figure S12 – Phylogenetic tree of complete haplogroup X3a sequences.** GenBank accessions and geographic origin are indicated for each complete sequence taken from the bibliography.

**Figure S13 – Phylogenetic tree of complete haplogroup U6a1a1 sequences.** GenBank accessions and geographic origin are indicated for each complete sequence taken from the bibliography.

**Figure S14 – Phylogenetic tree of complete haplogroup U6a7a1 sequences.** GenBank accessions and geographic origin are indicated for each complete sequence taken from the bibliography.

**Figure S15 – Phylogenetic tree of complete haplogroup U6b1a sequences.** GenBank accessions and geographic origin are indicated for each complete sequence taken from the bibliography.

**Figure S16 – Phylogenetic tree of complete haplogroup U6c sequences.** GenBank accessions and geographic origin are indicated for each complete sequence taken from the bibliography.

