## Supplementary figures and images for "Mitogenomes illuminate the origin and migration patterns of the indigenous people of the Canary Islands"

### Supplementary file 1

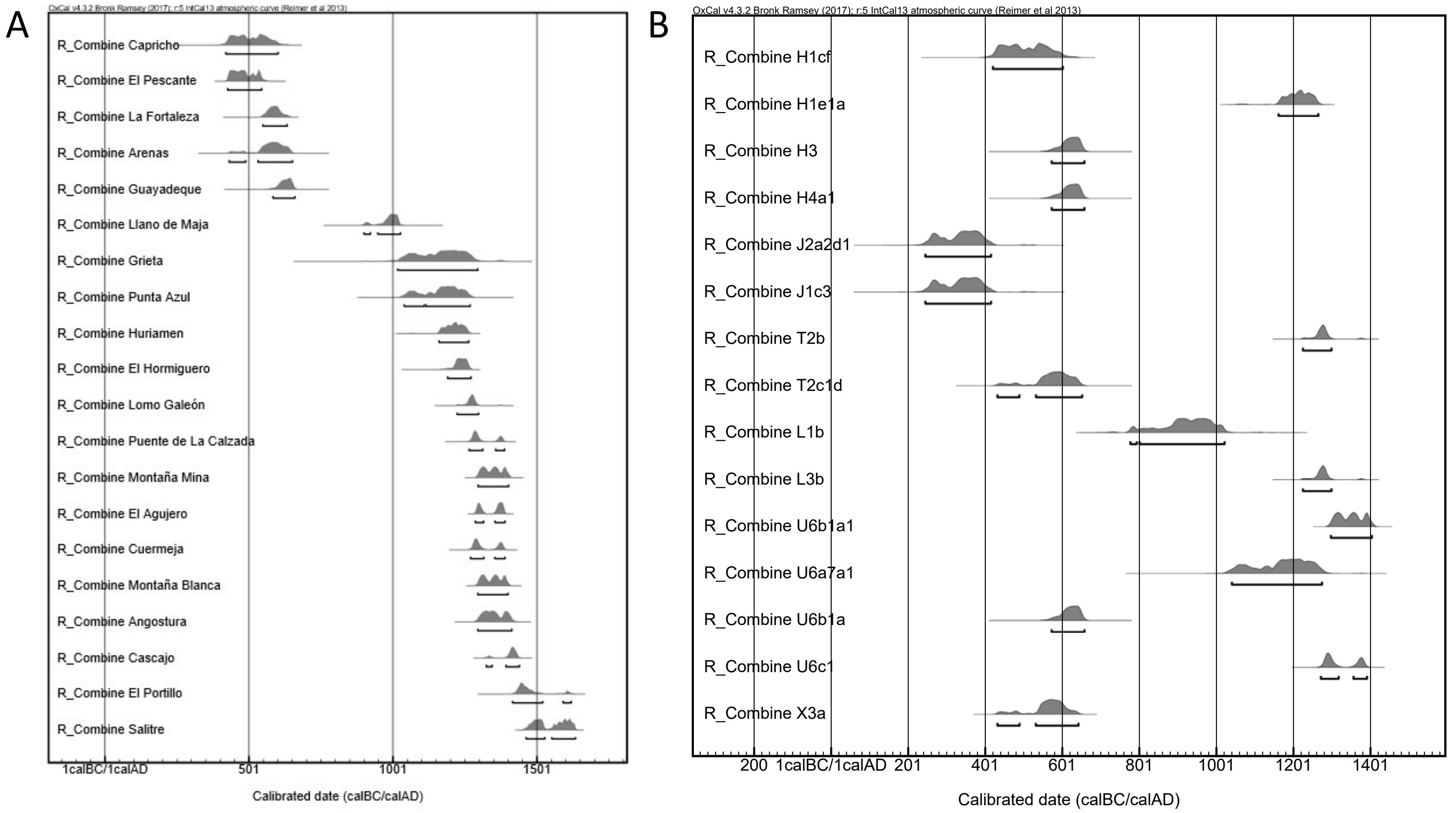

### Supplementary file 2

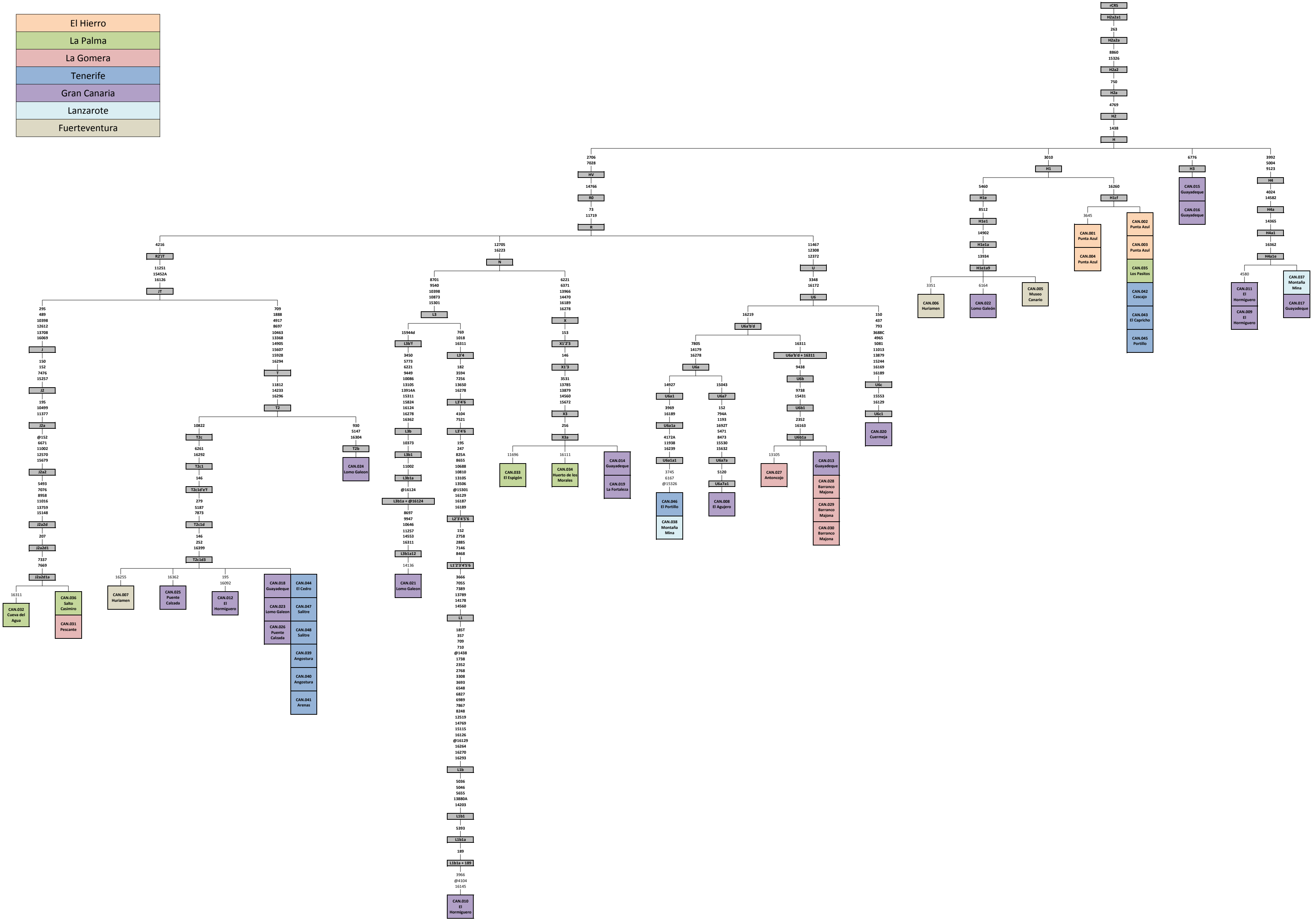

### Supplementary file 3

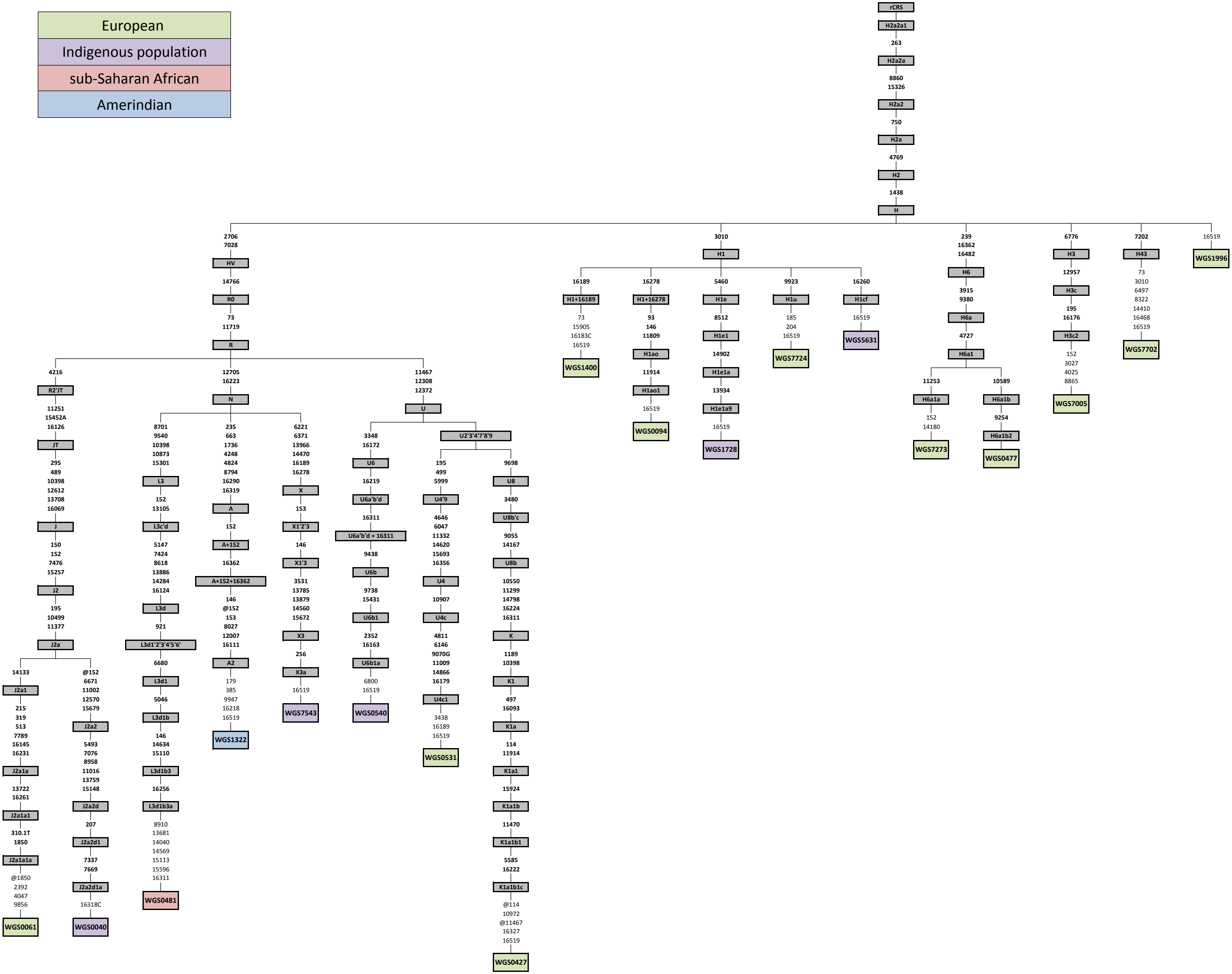

### Supplementary file 4

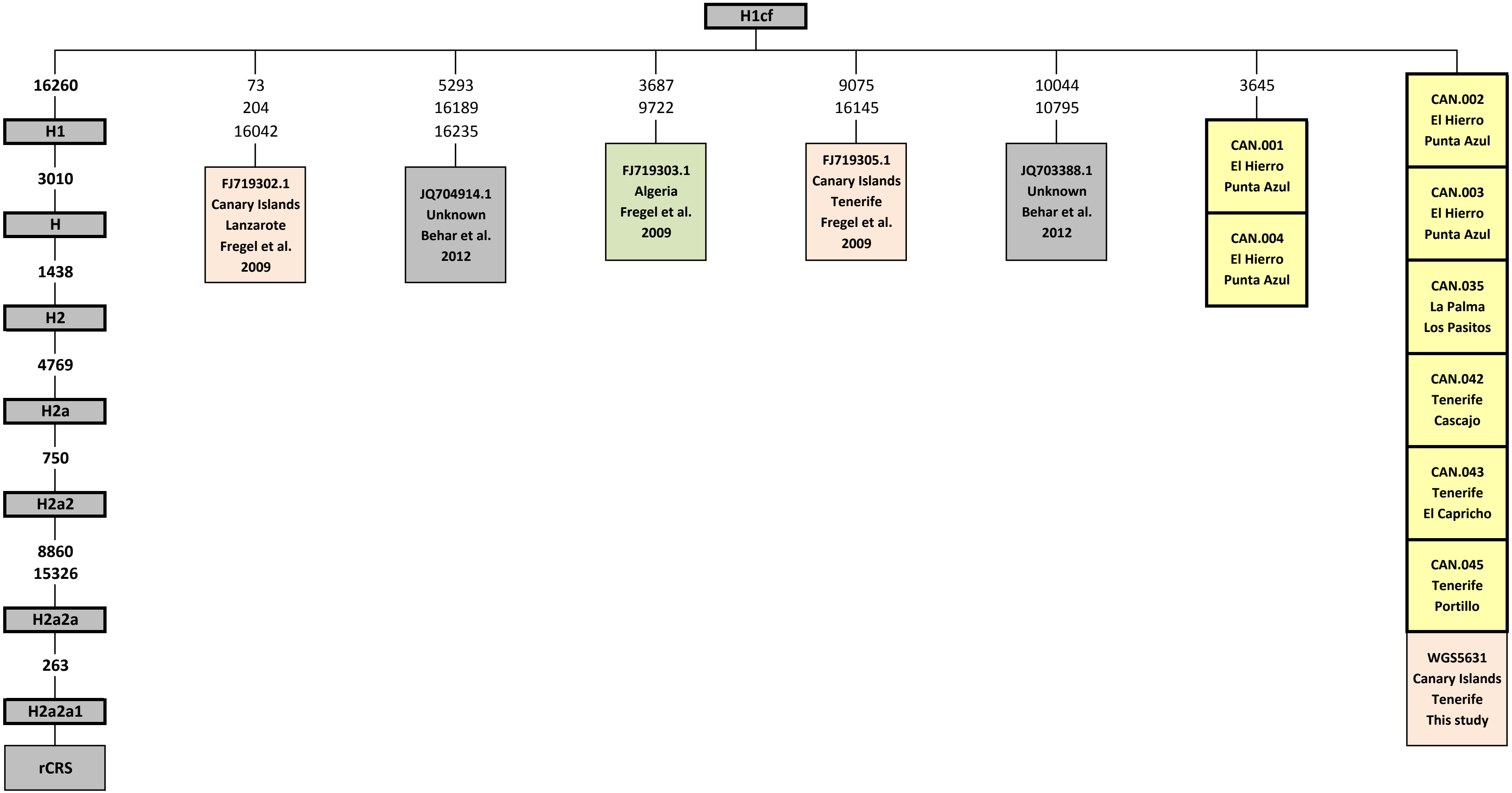

### Supplementary file 5

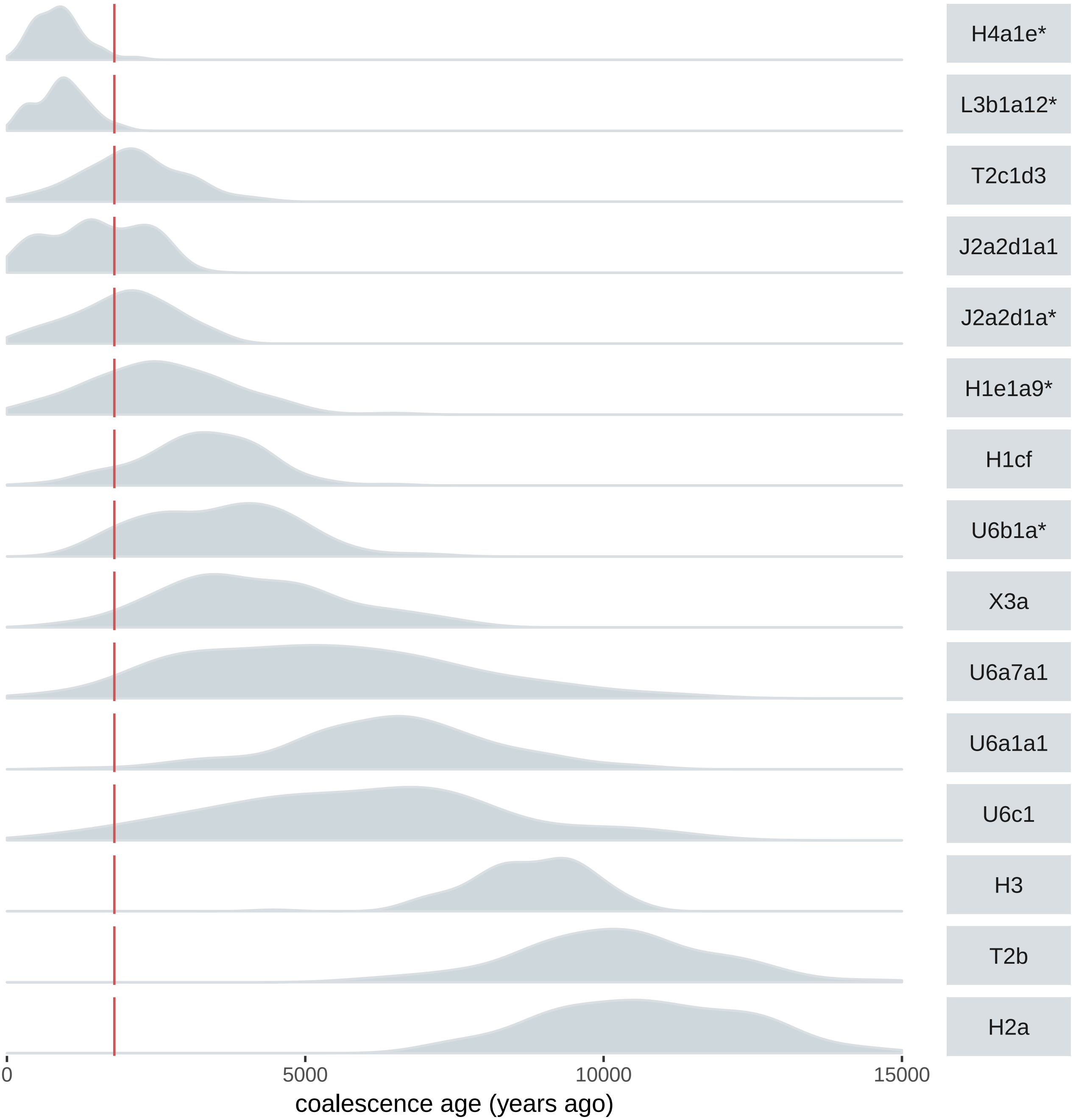

### Supplementary file 6

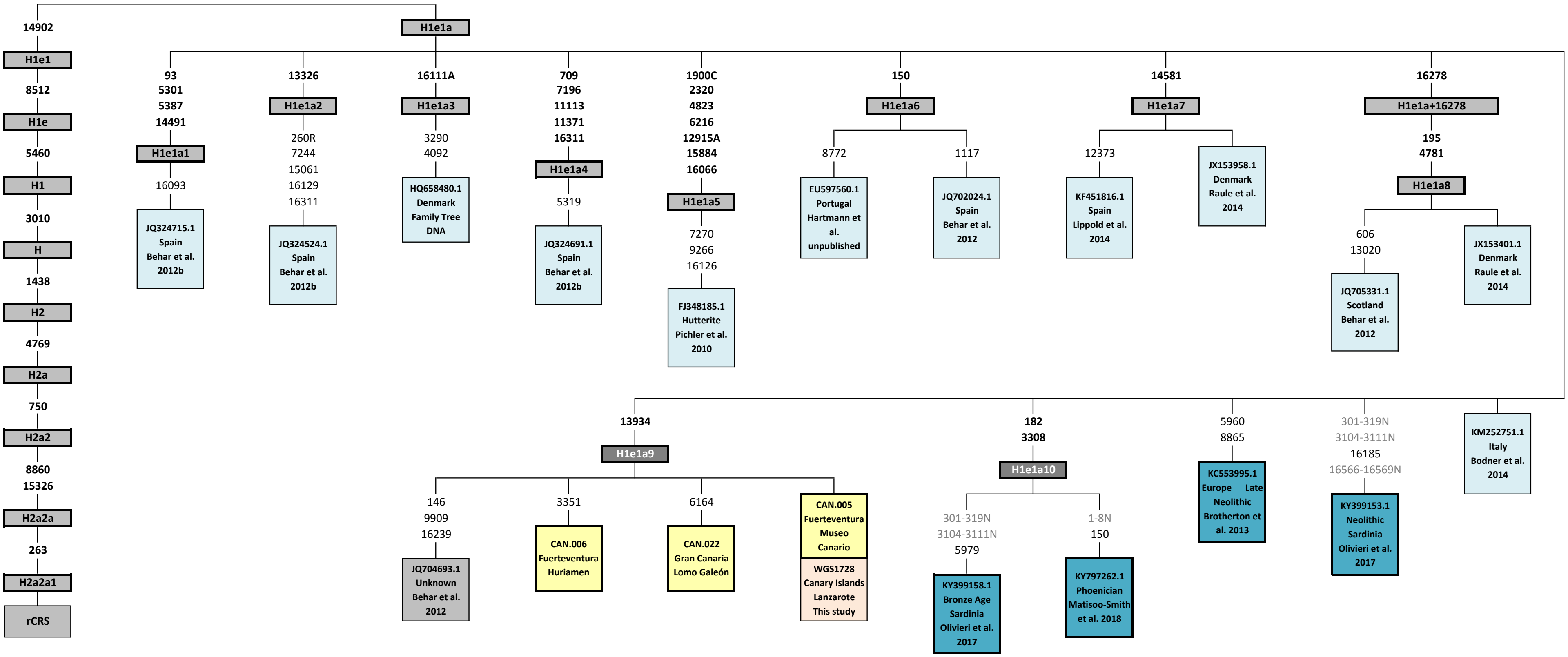

### Supplementary file 7

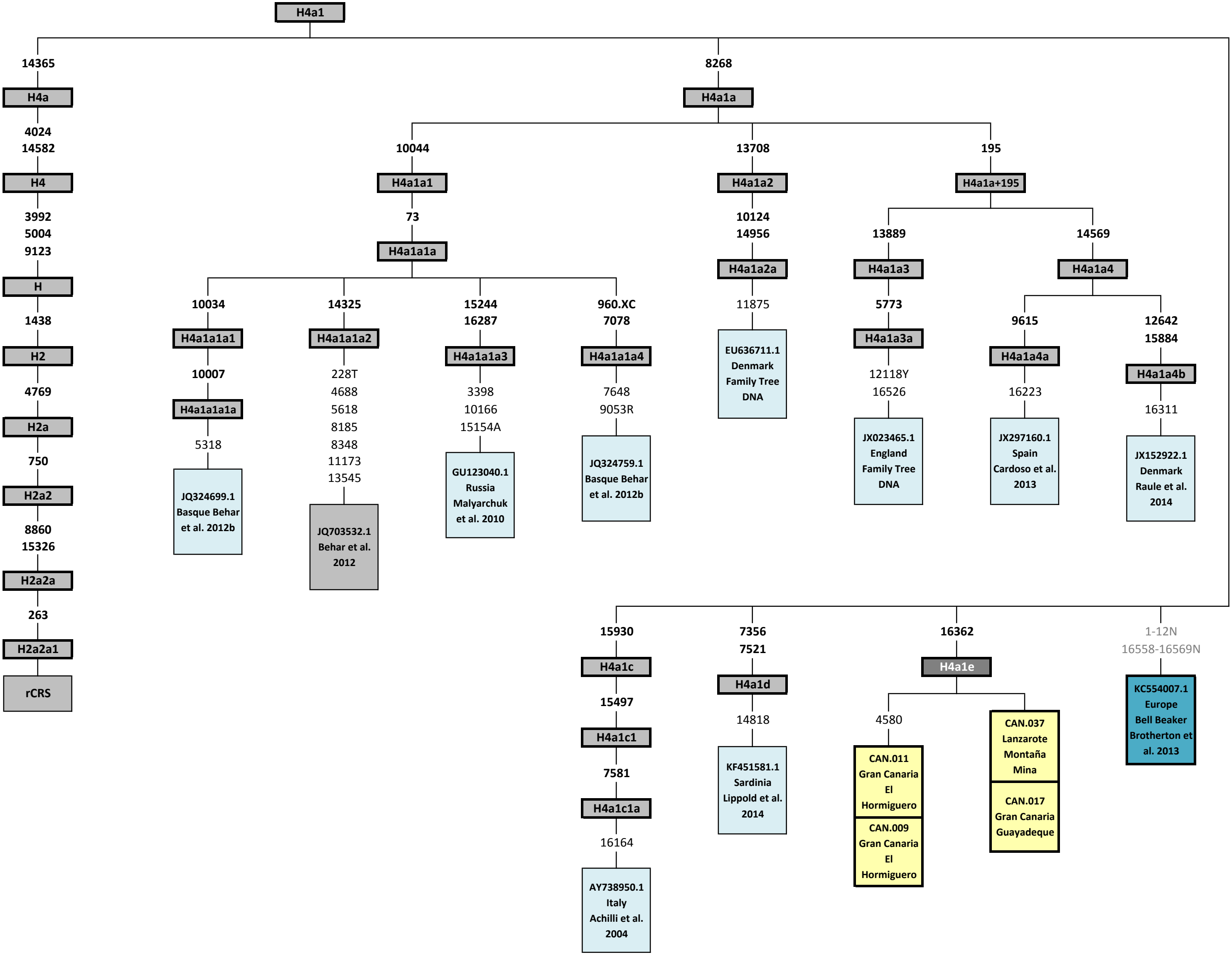

### Supplementary file 8

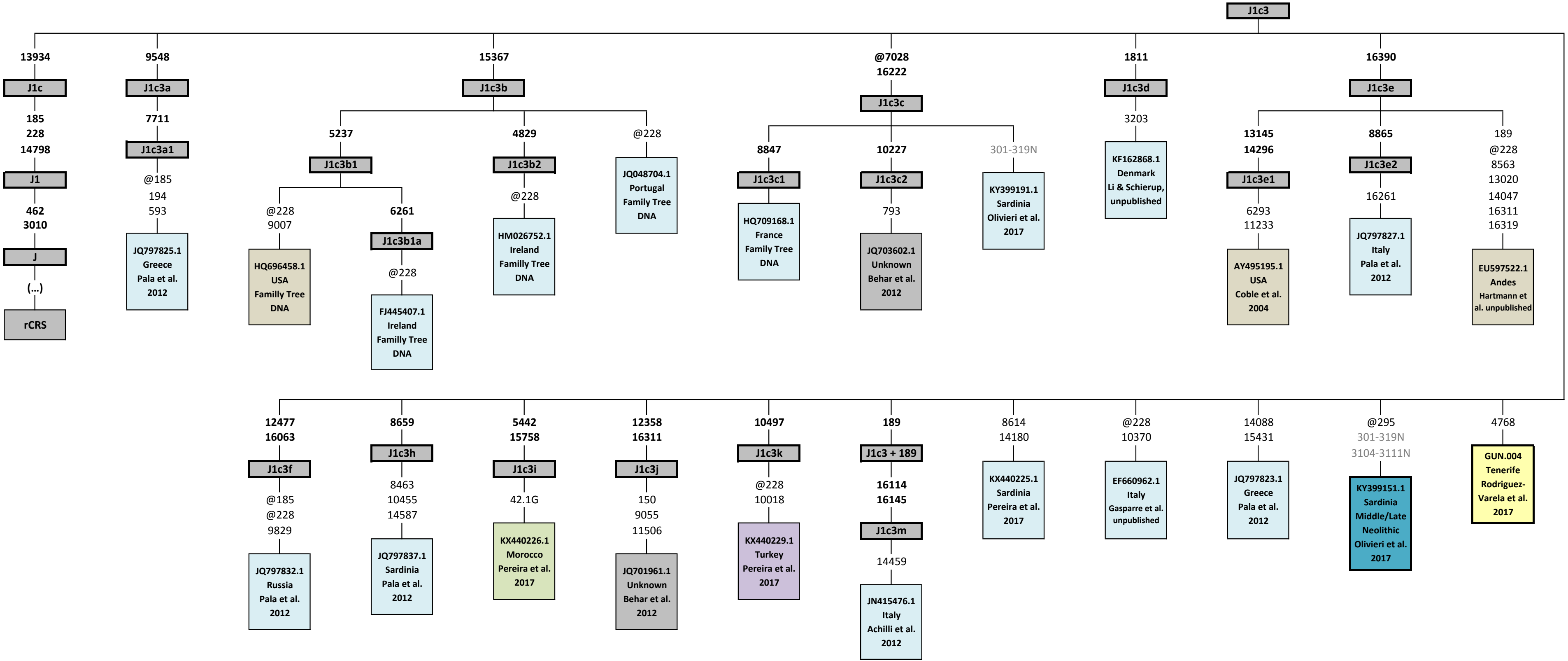

### Supplementary file 9

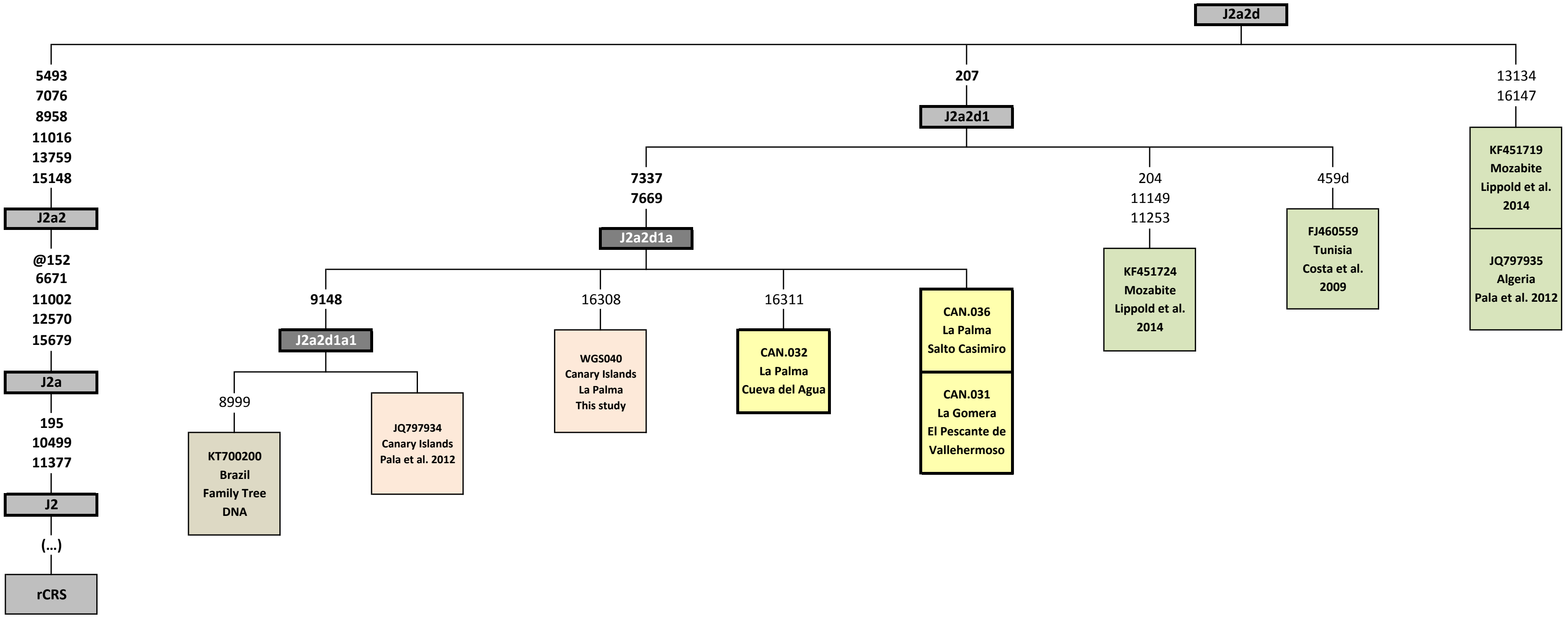

### Supplementary file 10

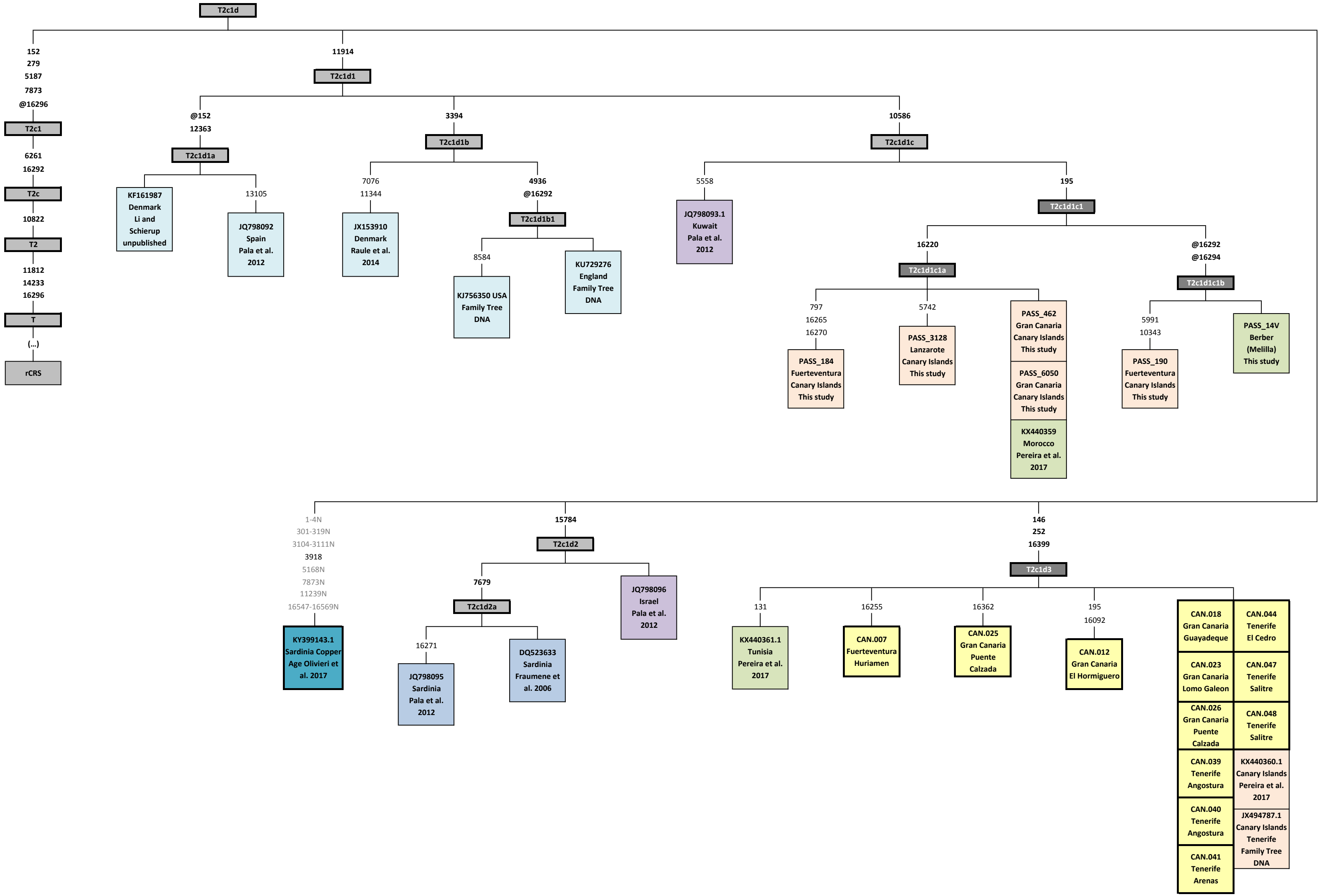

### Supplementary file 11

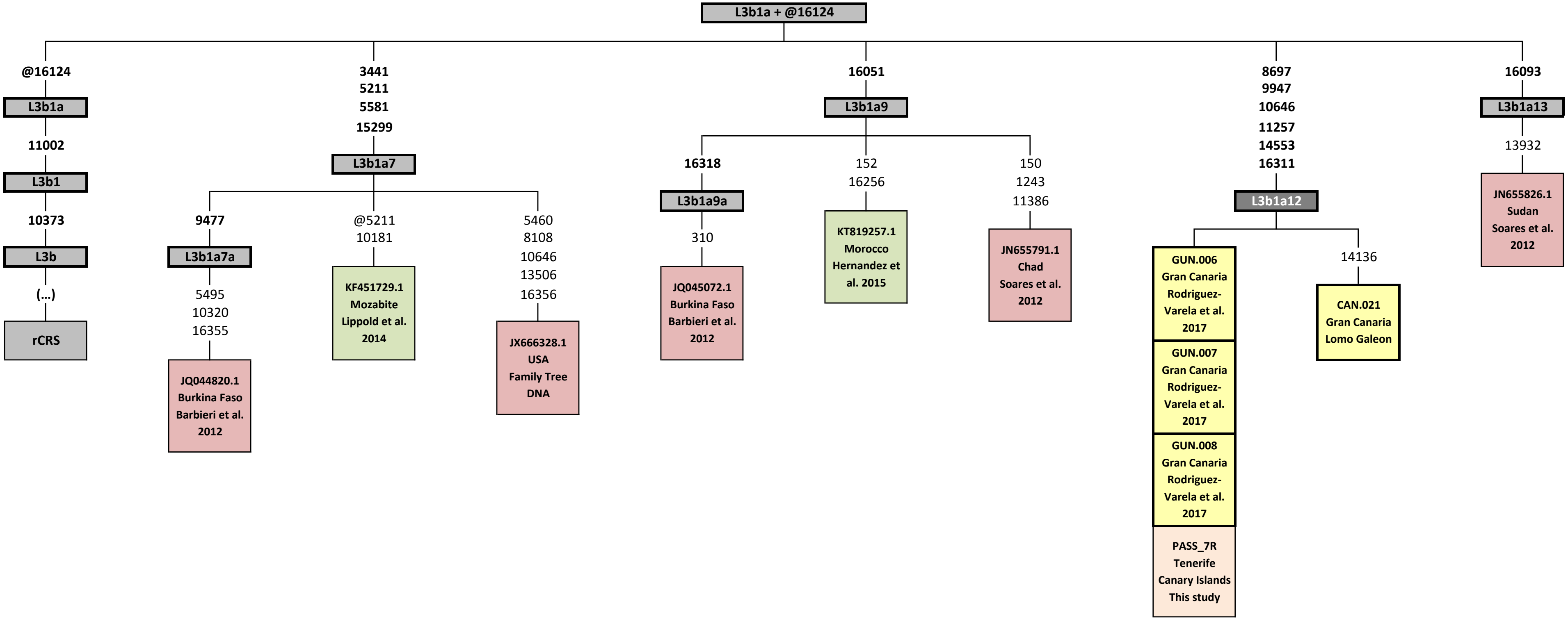

### Supplementary file 12

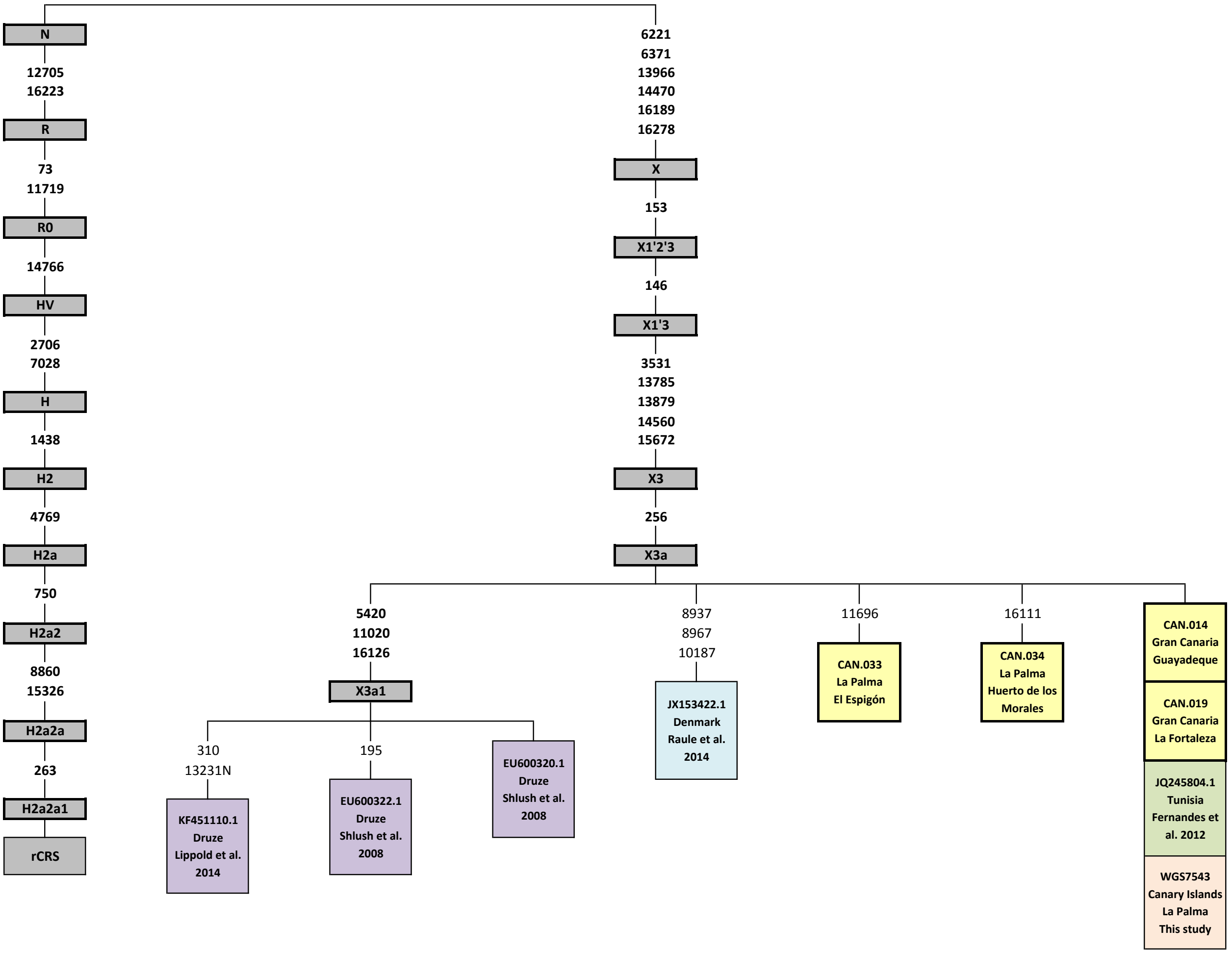

### Supplementary file 13

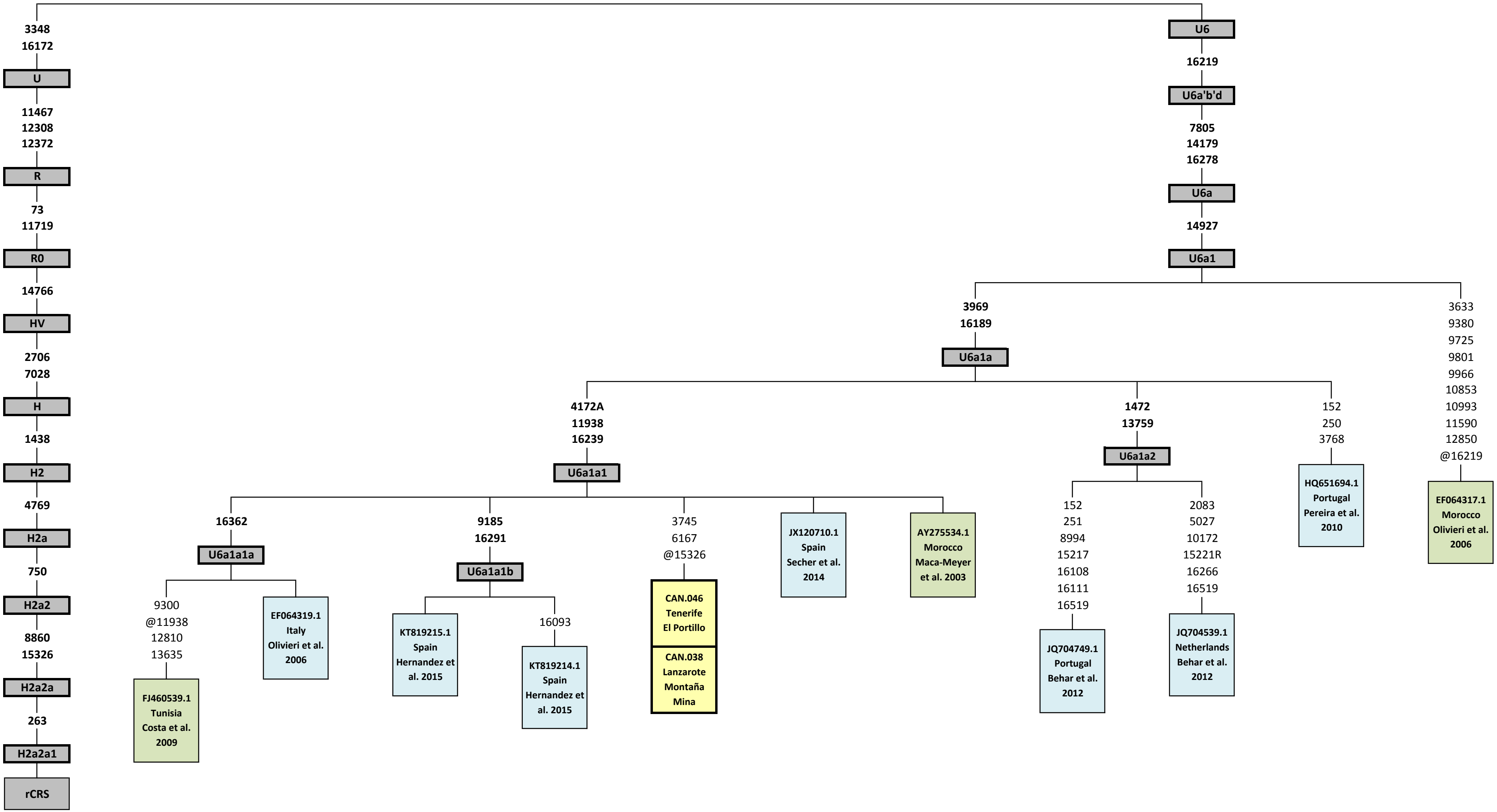

### Supplementary file 14

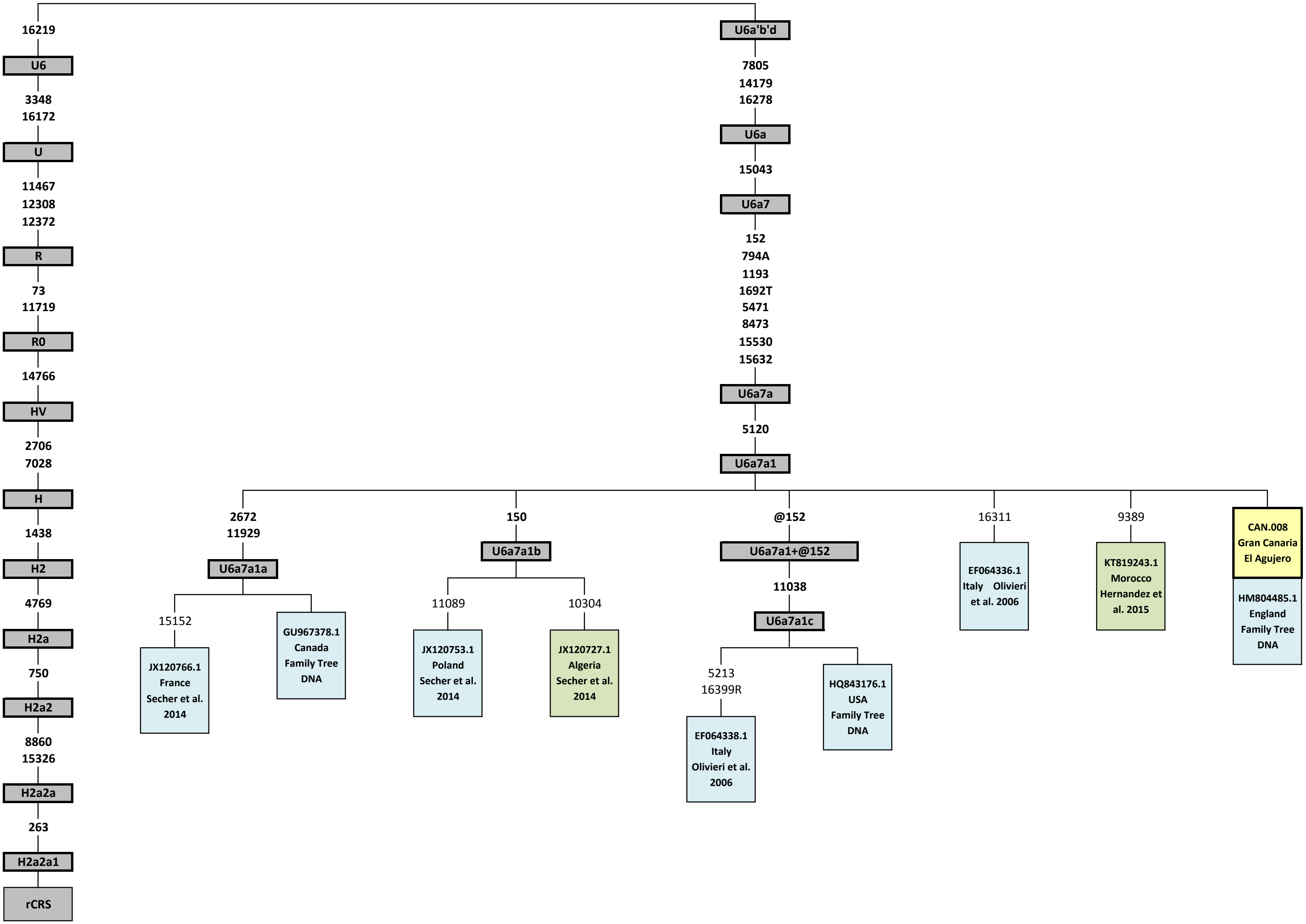

### Supplementary file 15

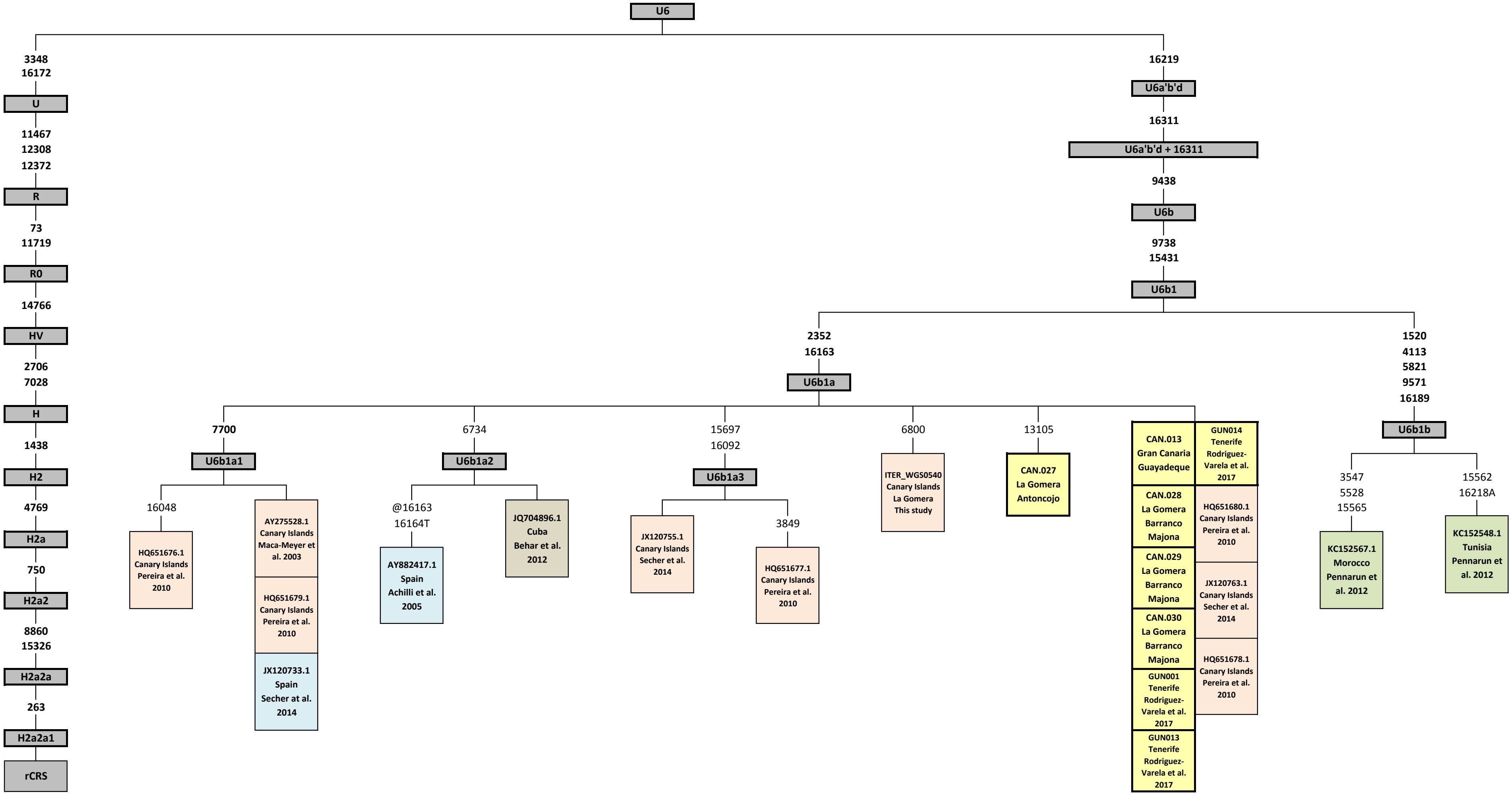

### Supplementary file 16

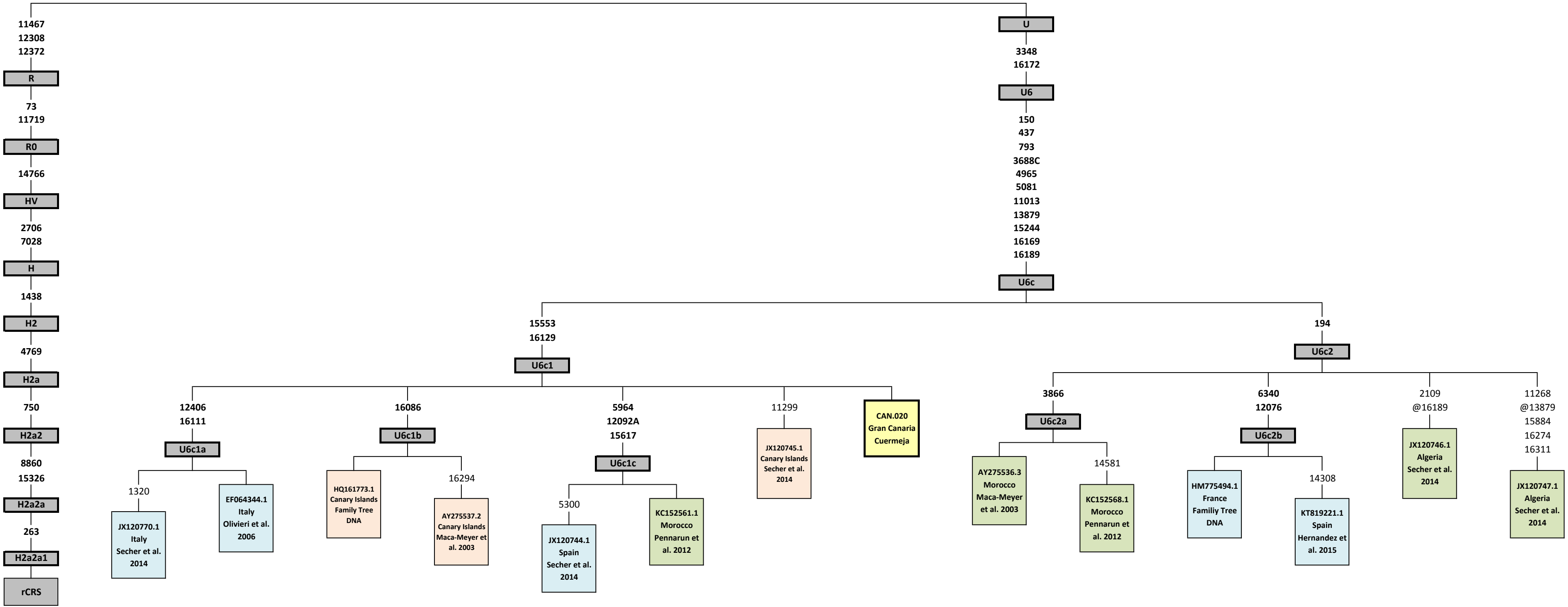
