## Supplementary material for "Mitogenomes illuminate the origin and migration patterns of the indigenous people of the Canary Islands"

Rosa Fregel, Departamento de Genética, Universidad de La Laguna. Avda. Astrofísico Fco. Sánchez, SN. San Cristóbal de La Laguna 38206, Spain.

### Detailed phylogeographic analysis of mitogenomes

#### Haplogroup H

Haplogroup H is the most frequent clade in the Iberian Peninsula and North Africa [1]. Frequencies of haplogroup H in North Africa increase westwards, being more frequent in Moroccans (~35%) than in Tunisians (~25%). The same H sub-haplogroups that are predominant in the Iberian Peninsula, such as H1 and H3, are also the most frequent in northwest Africa. On the other hand, Near Eastern clades like H4, H5, H7, H8 and H11 are more frequent eastwards. This asymmetrical distribution of H lineages in North Africa is believed to be related to a higher West European influence in the northwest and a higher Near Eastern impact in the northeast [1].

Haplogroup H is also the most frequent clade in our ancient sample, accounting for 36% of the population (Table S2). Two different H1 subhaplogroups are observed in the Canarian indigenous people: H1cf and H1e1a. Although haplogroup H1 is distributed across the entire European continent, it is especially common in West Europe. As we mentioned before, the highest frequency of H1 in Europe is observed in the Iberian Peninsula (18% - 24%). Given the distribution and coalescence age of H1 (~11,000 YBP), this haplogroup has been related to the late-glacial expansions of hunter-gatherers from the Franco-Cantabrian refuge [2], although this hypothesis has been challenged [3]. H1 frequency is relatively high in the Maghreb area (10% - 20%), reaching its peak in Tuaregs from Libya (61%). The high incidence of H1 in Northwest Africa has been presented as the result of Paleolithic expansions from the Iberian Peninsula into North Africa [1].

Based on complete mtDNA genome data, nine archaeological samples from the islands of El Hierro (Punta Azul), La Palma (Los Pasitos) and Tenerife (Cascajo, El Capricho and El Portillo) are classified as H1cf haplogroup, an H1 clade defined by 16260 (Fig S4). This confirms previously published HVRI data, which detected the presence of an H1-16260 haplotype in the indigenous people of Tenerife [4], La Palma [5], La Gomera [6] and El Hierro [7], and NGS data on one sample from Tenerife [8]. Apart from the ancient Canarian genomes, H1cf is observed in one sample from Algeria [5], and three from modern Canarian islanders (Lanzarote and Tenerife). In an HVRI analysis, a sample with a 16260 haplotype and belonging to the H1 haplogroup was also spotted in Tunisia [1]. Although haplogroup H1 is considered a West Eurasian lineage, based on our phylogeographic analysis (Fig S4), H1cf seems to be confined to both the Canary Islands and Central North Africa. Coincidently, the coalescence age of this clade is ~3,400 years (4,040 - 2,703 years ago), preceding the proposed time for the colonization of the islands (Fig S5).

Haplogroup H1e has a coalescence age of ~8,000 years [9]. Based on previous aDNA data from Neolithic archaeological sites, H1e was present in Europe since ~7,000 years ago [10]. H1e has also been related to current Jewish communities in Morocco [11]. Following the most updated version of phylotree (build 17), three archaeological samples from Fuerteventura (El Huriamen) and Gran Canaria (Lomo Galeón) are classified as H1e1a*. In our updated phylogenetic tree (Fig S6), H1e1a is observed mainly in modern European populations. In fact, a Neolithic sample from the Noeddale site Sardinia [12] and one ancient sample associated to the Late Neolithic Baalberge culture (~3,700 BC) [10] are also classified within H1e1* and H1e1a*, respectively. The presence of this haplogroup in the Canarian indigenous population would be in agreement with a European Neolithic expansion in North Africa [13]. All the ancient H1e1a samples share 13934 mutation, defining the new H1e1a9 subhaplogroup. One of our modern samples from Lanzarote also shares 13934 mutation, indicating that this lineage is still present in the modern population. However, discerning the distribution of this lineage in larger HVRI databases is impossible because of the lack of haplogroup-defining mutations in the control region and the scarcity of complete sequences in North Africa. Based on the information we have by now, this lineage can be considered exclusive of the Canary Islands. Unlike the autochthonous U6b1a, whose coalescence age is prior to the older radiocarbon dates associated to human presence in the

Canary Islands, the confidence interval of the coalescence age of H1e1a9 (~2,600 years ago) is compatible with an origin in the islands (Fig S5).

One sample from Gran Canaria from Rodriguez-Varela et al. [8] is classified within H2a, with two private mutations (3010 and 9053) that have not been observed in any of the H2a sequences available in the NCBI database. While H1 and H3 haplogroups have their highest frequencies in Western Europe, H2 is more abundant in Eastern Europe. Haplogroup H2a has also been observed in North Africa [1], but its frequency is lower than that of H1 (1.3%).

Two archaeological samples from Gran Canaria (Guayadeque) are classified as H3*, with no private mutations, so preventing further phylogeographic analysis. Haplogroup H3 is found in the Maghreb area, with frequencies around 1.5 - 3.5%. Coalescence age and distribution of H3 is similar to those of H1, and it is considered to be related to Paleolithic expansions from Europe into North Africa [1].

Three samples from Gran Canaria (El Hormiguero and Guayadeque) and one from Lanzarote (Montaña Mina) are classified as H4a1, one of the linages that are restricted to the eastern islands. Haplogroup H4 is present mostly in the Near East [14], although it appears at a low frequency in Europe [15] and North Africa [1]. In fact, the presence of H4 in North Africa has been associated with gene flow from the Near East. In our phylogeographic analysis, H4a1 sublineage is observed only in Europe and the Canary Islands (Fig S7). All ancient samples share mutation 16362, defining the new H4a1e haplogroup. As haplogroup H4a1 is also observed in one Bell Beaker sample from Europe, its presence in the Canarian indigenous people, and more generally in North Africa, could be related to a European Chalcolithic intrusion rather than a Near Eastern influence. Regarding the distribution of the newly defined H4a1e, in a HVRI database with H samples from the Iberian Peninsula (n=480), North Africa (n=880), the Arabian Peninsula (n=1,115) and the Near East (n=152) [1], no sample classified within H4 by RFLP analysis showed the 16362 mutation in the control region, indicating that this lineage may have a similar distribution pattern as U6b1a. In this case, the coalescence age for H4a1e (~860 years ago) overlaps with the period of human occupation of the Canary Islands (Fig S5).

#### Haplogroup J

Four ancient samples from the Canary Islands belong to haplogroup J. One sample from Tenerife (archaeological site unknown) [8] is classified within J1c3, and one sample from La Gomera (Pescante) and two from La Palma (Salto del Casimiro and Cueva del Agua) are clustered within J2a2d1 haplogroup. Haplogroup J is relatively frequent in both the Near East (~13%) and Europe (9%), although a phylogeographic analysis supports that this clade initially diversified in the Near East around 33,000 years ago [16]. Haplogroup J1 represents the vast majority of the clade's lineages. Although J1a and J1b are of Near Eastern distribution, J1c is considered to be a European cluster. The ancient sample from Tenerife [8] is classified as J1c3 with a private mutation at position 4768 that was not found in any J1c sequence from the GenBank database (14/11/2017), thus remaining as J1c3* (Fig S8). This haplogroup and its sublineages have been observed in Europe, North Africa and the Near East. Interestingly, J1c3 has also been observed in a Neolithic sample from Spain belonging to the Cardial culture [17] and a Middle/Late Neolithic samples from Sardinia [12]. As both paleogenomic and archaeological records indicate the existence of contact and exchange between Iberia and the Maghreb [13,18,19], the presence of this lineage in the indigenous people of the Canary Islands could be explained again by prehistoric migrations into North Africa.

Haplogroup J2 is less frequent than J1 and is mostly related to the Near Eastern, although some subclusters are of European affiliation. However, J2a2d is restricted to Central North Africa (Algeria and Tunisia) and the Canary Islands [16]. In our phylogeographic analysis, we observe that one Brazilian sample is also classified within the J2a2d clade (Fig S9). However, this result can be explained by historical migrations of Canarians to Latin America [20]. Accordingly, all the Canarians and the Brazilian samples share mutations 7337 and 7669, defining the new J2a2d1a sub-haplogroup. In addition, one Canarian and the Brazilian sample share mutation 9148, defining the J2a2d1a1 sub-lineage. Based on these results, the Canarian clade also qualifies for being considered an autochthonous lineage of the islands. This is tricky because of the lack of mutations in the control region and the impossibility of performing a search on the larger HVRI datasets. Nevertheless, the coalescence age of the clades J2a2d1a (~2,151 years ago) and J2a2d1a1 (~1,300 years ago) could be in agreement with an origin in the Canary Islands (Fig S5).

#### Haplogroup T2

One sample from Gran Canaria (Lomo Galeón) is T2b, while a total of 13 samples from several archeological sites in Tenerife, Gran Canaria and Fuerteventura are T2c1d. Haplogroup T2 most likely originated in the Near East ~20,000 years ago [16]. Today, T2 is frequent in the Mediterranean and Western Europe but is also common in the Near East. T2b is considered to have originated ~10,000 years ago in Europe and dispersed during the early Neolithic period. Numerous aDNA Neolithic samples are classified within T2b, including archaeological sites from France, Germany, Italy, Sweden and Spain. It is worth mentioning that one Late Neolithic sample from Morocco has been assigned to haplogroup T2b3, indicating the presence of T2b in North Africa since 3,000 BCE [13]. The sample from Gran Canaria analyzed in this study is classified as T2b, with no private mutations, precluding further phylogenetic analyses.

T2c originated in the Near Easter ~20,000 years ago and later expanded into Europe. Most of the T2c lineages belong to haplogroup T2c1, which also appears to have a Near Eastern origin ~18.500 years ago and spread into Europe ~10,000 years ago [16]. T2c1 is frequent in Cyprus and the Persian Gulf, but it is also present in the Levant and the Mediterranean regions. In our phylogenetic tree (Fig S10), haplogroup T2c1d appears distributed in Europe, the Near East, North Africa and the Canary Islands. It is worth mentioning that T2c1d lineages have been observed in several European Neolithic and Bronze Age sites [13,21,22]. In fact, a Bronze Age sample from Sardinia is classified within T2c1d* [12]. All the Canarian ancient samples, two samples from the modern Canarian population and one sample from Tunisia [23] are classified as T2c1d* based on the last version of the phylotree [24], but share mutations 146, 252 and 16399, defining haplogroup T2c1d3. As this haplogroup is found both in ancient samples from the Canary Islands and Tunisia, it might connect the indigenous population with Central North Africa.

As commented earlier, mtDNA analysis on the modern population of the Canary Islands suggested that the asymmetrical distribution of T2c1 could be evidence of a second indigenous migration wave. The same asymmetrical distribution is observed for aDNA data in this study when the entire dataset is used for calculations. The coalescence age for T2c1d3 is relatively recent, ~2,150 years ago (2,858 - 1,431) (Fig S5). If we compare mtDNA results with radiocarbon dates reported for each archaeological site (Fig. S1), the earlier dates for T2c1d3 observed in the archipelago are in one sample from the Guayadeque site in Gran Canaria (540 to 737 AD), one sample from Las Arenas in Tenerife (540 to 650 AD) and in one decontextualized sample from Tenerife (792 AD) [8]. Taking into account that the oldest calibrated radiocarbon dates are from the $3^{rd}$ - $5^{th}$ century, T2c1d3 seems to have been present in the two central islands at relatively early stages.

Interestingly, the modern DNA samples included in this study for their classification as T2c1 based on HVRI data (five samples from the Canary Islands and two from North Africa) fall within T2c1d1 and share mutation 10586 with a sample from Kuwait [16], defining the new T2c1d1c haplogroup (Fig S10). All the newly sequenced samples share mutation 195 and define haplogroup T2c1d1c1. Additionally, four modern Canarian samples and one Moroccan sample share mutation 16220 (T2c1d1c1a), and one Canarian sample and one Berber sample from Melilla share @16292 and @16294 mutations (T2c1d1c1b). It is worth mentioning that all the Canarian samples classified within the new T2c1d1c1 have been sampled in the eastern islands (Gran Canaria, Lanzarote and Fuerteventura). If we analyze the distribution of the control region haplotype (16126 16220 16292 16294) in the larger HVRI database of the Canary Islands (n=803) [25], T2c1d1c1a is confirmed to be restricted to the eastern islands and differences between eastern/western populations are significant ($p < 0.0001$). Although the presence of this lineage has not been confirmed in pre-colonial times, it might reflex additional prove of the asymmetrical distribution of maternal lineages in the islands. However, this distribution could also be related to the later migration of "moriscos" (Moorish slaves) to the islands in colonial times [26,27]. Giving that the impact of the Moorish slave trade was higher in the eastern islands and that T2c1d1c1 has not been observed in the indigenous population, this last explanation seems more plausible.

##### Haplogroup L1b

L1b is a sub-Saharan African lineage that is more frequent in West Africa [28]. L1b is observed in North Africa, especially in Morocco (6.1%) and West Sahara (11.4%) [29]. One ancient sample from Gran Canaria (El Hormiguero) is classified as L1b1a+189 with several private mutations (3966, @4104 and 16145) that have not been observed in any of the complete L1b sequences in the GenBank database. Based on the coalescence ages of certain L1b lineages in North Africa, and more generally the Mediterranean region, prehistoric sub-Saharan African migrations have been proposed to have reached those areas around the Holocene Climatic Optimum (9,000 to 5,000 years BP) [30]. Although Later Stone Age (~15,000 BP) [31] and Early (~7,000 BP) and Late Neolithic (~5,000 BP) samples from North Africa did not show any mtDNA lineage of sub-Saharan origin [13], our results indicate the presence of L1b in North Africa at least at the time of the colonization of the Canary Islands. It is worth mentioning that this particular haplotype is absent in the HVRI dataset of the modern population of the Canary Islands [25].

##### Haplogroup L3b

L3b is also a sub-Saharan mtDNA lineage [32] that most probably originated in Central/West Africa [33]. One of our ancient samples from Gran Canaria (Lomo Galeón), three samples from Gran Canaria from Rodriguez-Varela et al. [8] and one modern sample from Tenerife fall within L3b1a+@16124. Interestingly, based on HVRI data, haplogroup L3b1a+@16124 has been spotted at low frequencies in North Africa, particularly in Andalusians from Tunisia (1.96%) and Libya (1.86%) [29]. In our summarized phylogenetic tree (Fig S11), L3b1a+@16124 is present both in sub-Saharan and North Africa. Also, all the samples from the Canary Islands share six private mutations (8697 9947 10646

11257 14553 16311) and define the new L3b1a12. By looking at the easily recognizable motif of L3b1a12 (16223 16278 16311 16362) in a large HVRI database, this haplogroup is observed today in Tenerife, La Gomera and Gran Canaria, with an overall frequency in the archipelago of 1.0%. Interestingly, although this haplogroup has not been observed in the ancient population of Tenerife, today is as frequent there (2.03%) as in Gran Canaria (1.41%). Although this result may be associated to sampling issues, it is suggestive of the historical chronicles of indigenous people of Gran Canarian aiding the Castilians during the conquest of Tenerife, and later being rewarded with land ownership in the island [34]. The recent coalescence age of L3b1a12 (~860 years) is in agreement with an origin in the archipelago (Fig S5).

**Haplogroup X3**

Haplogroup X3 has a Mediterranean distribution and it is considered to be restricted to the Near East and northeast Africa [35]. Two ancient samples from Gran Canaria (Guayadeque and La Fortaleza) and two from La Palma (El Espigón and Huerto de Los Morales), and one modern sample from La Palma are classified as X3a. Although both ancient samples from La Palma present private mutations, they have not been observed in any of the publicly available complete sequences from GenBank. In our phylogenetic tree (Fig S12), we confirm the presence of X3a in Europe, the Near East and North Africa, concretely in Tunisia. The coalescence age for this lineage (~4,200 years ago) is consistent with a Chalcolithic expansion in the Mediterranean (Fig S5).

**Haplogroup U6**

U6 is considered a North African autochthonous haplogroup associated to the back migration to Africa from Eurasia during the Paleolithic [36,37]. In fact, the presence of haplogroup U6 in relative frequencies in modern-day North African has been presented as evidence of maternal continuity in the region since Paleolithic times. However, although U6 is autochthonous in North Africa, its phylogeny is better explained admitting both prehistoric and historic U6 influences in Europe [37]. One sample from Tenerife (El Portillo) and one from

Lanzarote (Montaña Mina) belong to U6a1a1 haplogroup. Both samples have three private mutations but none of them have been observed in the GenBank database. Our phylogenetic tree (Fig. S13) indicates that U6a1a1 is present in the Maghreb, South Europe and the indigenous people of the Canary Islands.

One sample from Gran Canaria (El Agujero) is classified as U6a7a1. This haplogroup is considered to be a U6 lineage with a European distribution, which most probably spread within the Chalcolithic period (~4,700 years ago) [37]. In our phylogeographic analysis, U6a7a1 is prominently observed in European populations, but also spotted in Morocco, Tunisia and Algeria (Fig. S14).

Four samples from La Gomera (Barranco Majona and Antoncojo), one from Gran Canaria (Guayadeque) and three samples from Tenerife [8] are classified as U6b1a. As we mentioned before, this haplogroup has been of interest as it is considered to be autochthonous in the Canary Islands, as its presence in Cuba and Spain are attributed to historical migrations. However, the coalescence age of ~3,600 years ago predates the proposed time for the colonization of the islands (Fig S5). On the other hand, the three sub-haplogroups within U6b1a better coincide with the timeline based on radiocarbon dating: U6b1a1 (~1,500 years ago), U6b1a2 (~2,600 years ago) and U6b1a3 (~1,300 years ago). This result indicated that U6b1a most probably originated in North Africa and later migrated to the Canary Islands, where it diversified [37]. The absence of U6b1a in modern North Africa could be related to later population events modifying the mtDNA composition. Interestingly, all the ancient U6b1a lineages but one have the basal motif, and the remaining one is also classified as U6b1a* with a private mutation not observed in the modern population (Fig S15).

U6c1 was previously considered a Canarian autochthonous lineage but was later associated with a Mediterranean distribution, by being discovered in southern Italy and Tunisia [37]. One ancient sample from Gran Canaria belongs to U6c1* with no private mutations (Fig S16). Another modern sample from the Canary Islands is classified as U6c1*, with one private mutation. Sub-haplogroups within U6c1 have clear geographical adscriptions: U6c1a is restricted to Italy, U6c1b to the Canary Islands and U6c1c is observed both in Spain and Morocco [37]. The coalescence age for the whole U6c1 haplogroup is ~6,000 years, which is congruent with an origin outside the islands; however, the Canarian-specific U6c1b is 1,300 years and most probably originated in the archipelago.
